# Are under-studied proteins under-represented? How to fairly evaluate link prediction algorithms in network biology

**DOI:** 10.1101/2022.10.13.511953

**Authors:** Serhan Yılmaz, Kaan Yorgancioglu, Mehmet Koyutürk

**Affiliations:** Department of Computer and Data Sciences, Case Western Reserve University; Center for Proteomics and Bioinformatics, Case Western Reserve University

## Abstract

For biomedical applications, new link prediction algorithms are continuously being developed and these algorithms are typically evaluated computationally, using test sets generated by sampling the edges uniformly at random. However, as we demonstrate, this evaluation approach introduces a bias towards “rich nodes”, i.e., those with higher degrees in the network. More concerningly, this bias persists even when different network snapshots are used for evaluation, as recommended in the machine learning community. This creates a cycle in research where newly developed algorithms generate more knowledge on well-studied biological entities while under-studied entities are commonly overlooked. To overcome this issue, we propose a weighted validation setting specifically focusing on under-studied entities and present AWARE strategies to facilitate bias-aware training and evaluation of link prediction algorithms. These strategies can help researchers gain better insights from computational evaluations and promote the development of new algorithms focusing on novel findings and under-studied proteins.

**Teaser:** Systematically characterizes and mitigates bias toward well-studied proteins in the evaluation pipeline for machine learning.

**Code and data availability:** All materials (code and data) to reproduce the analyses and figures in the paper is available in figshare (doi:10.6084/m9.figshare.21330429). The code for the evaluation framework implementing the proposed strategies is available at github^†^. We provide a web tool^‡^ to assess the bias in benchmarking data and to generate bias-adjusted test sets.

## Introduction

### Background and related literature

Link prediction (LP) is a widely used technique in network biology for discovering previously unknown associations or interactions among biological entities [1–9]. Early research in this field focused on local measures such as common neighbors and preferential attachment [10, 11], in addition to global approaches such as random walks that consider node proximity as an indicator of potential edges [12]. More recently, graph embedding models have emerged, allowing machine learning techniques to be seamlessly integrated into link prediction [13, 14]. With the availability of diverse omic data and advancements in machine learning, more sophisticated algorithms like graph convolutional networks are increasingly applied to link prediction in systems biology [15–17].

Link prediction algorithms face a severe class imbalance challenge [18]. Although conventional machine learning offers various approaches to address this issue, such as under-sampling [19, 20], over-sampling [21, 22], and their combinations [23], applying them to link prediction is challenging due to the quadratic growth of the search space with the number of nodes. As a result, the prevailing approach is to use simpler techniques like under-sampling possible node pairs uniformly at random [1, 13].

### Evaluation of link prediction algorithms

For evaluating link prediction algorithms, a recommended approach is to use an independent test dataset and train the algorithms on different snapshots of the network from various data sources or time points [18, 24]. In cases where multiple snapshots are not available, the evaluation is typically conducted by randomly sampling edges to be removed from a single network [14, 25, 26]. While there has been considerable attention given to algorithmic bias, fairness, reproducibility, and comparability in graph machine learning [27, 28], studies specifically investigating fairness and sources of bias in evaluating link prediction algorithms are relatively limited [24, 29], particularly in the context of network biology [30].

### Motivation and significance in systems biology

Fairness in biological knowledge discovery involves addressing the under-representation of less studied biological entities. However, the Matthew’s effect, commonly known as “rich gets richer”, is highly prevalent in biology. As highlighted by the Understudied Protein Initiative [31], “95% of all life science publications focus on a group of 5,000 particularly well-studied human proteins”. This pronounced bias poses a critical challenge during the evaluation of link prediction algorithms in biology. While previous studies have documented degree bias in biological networks and its implications in specific network biology applications [4, 30, 32], limited attention has been given to the impact of bias on evaluating new link prediction algorithms [33].

Consequently, algorithms that make biased predictions toward high-degree nodes, which often correspond to well-studied proteins [34], are disproportionately rewarded, as we show in our results. This creates a serious barrier to making new discoveries involving under-represented proteins, as the potential for developing specialized algorithms targeting under-studied proteins is overlooked. As a result, progress in these areas is hindered.

### Contributions of this study

Motivated by these considerations, using prediction of protein-protein interactions (PPI) as a case example, we investigate how typical evaluation settings in the literature incentivize algorithms to prioritize well-studied proteins in their predictions. To facilitate bias-aware evaluation, we propose multiple strategies to address various aspects of the issue. These include quantifying algorithmic bias, addressing bias in benchmarking data, implementing balanced evaluation settings with weighted metrics or stratified analysis to focus on under-studied proteins, and providing a comprehensive overview using diverse metrics for bias-aware comparisons and summarizing research findings. Additionally, we explore a degree-balanced under-sampling approach for training of link prediction algorithms to enhance predictions involving under-represented proteins.

## Results

### Experimental Setting

In this paper, our main goal is to highlight issues in standard evaluation settings arising from the severe imbalance in our knowledge of the biological networks, as well as to demonstrate the strategies we propose to resolve these issues. In our experiments, we mainly utilize the Biogrid protein-protein interaction network [35] as a case study. As a final part of our study, we extend our analysis to various networks and problems within network biology, demonstrating that our findings regarding PPI networks are generalizable across other biomedical domains as well. For simplicity, we discuss nodes primarily in a protein context, although the developed techniques are not limited to proteins or PPI network.

To select the link prediction algorithms for our analysis (Figure 1a), we consider two criteria. First, we aim to include representative methods from different algorithm classes, such as scoring metrics, network propagation methods, and embedding/machine learning methods. Second, we aim to include algorithms that exhibit varying levels of bias towards high-degree nodes. For each category, we include at least two versions: one high-bias version that prioritizes high-degree nodes, and another normalized version with lower bias. For example, in the scoring metrics category, the high-bias version is common neighbors, while the normalized, lower-bias version is the Jaccard index. Both versions utilize the same information source (number of shared interactions), but the normalization based on node degrees differs, influencing the algorithm’s inclination towards high-degree nodes.

**Figure 1:**
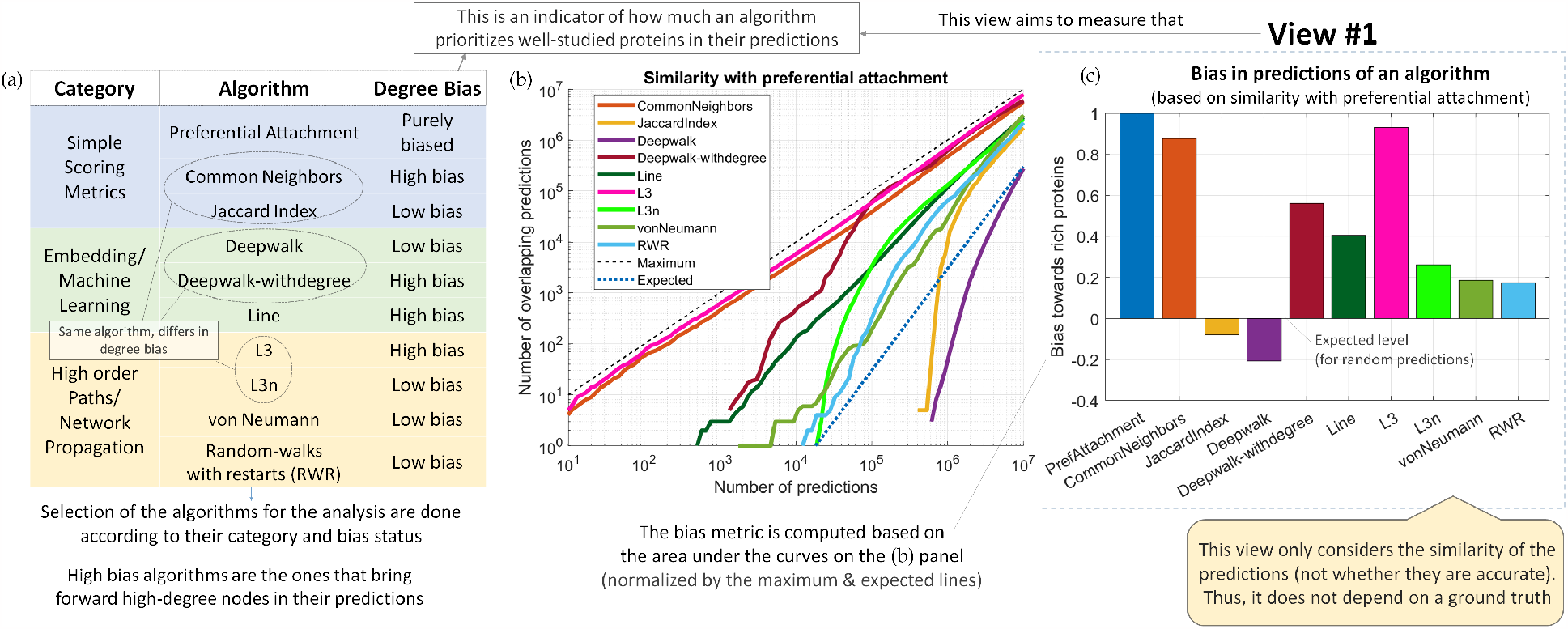
Quantifying the bias towards well-studied proteins in the predictions of an algorithm based on similarity with preferential attachment model on Biogrid PPI network. (a) The algorithms selected for the analysis, their categorization and affinities towards degree bias. (b) Overlap of the predictions of the algorithms with preferential attachment. (c) The quantified bias of the algorithms.

We also extend Deepwalk [13], a low-bias embedding algorithm that generates feature vectors with minimal correlation to node degrees (S. Figure 1). This biased variant, Deepwalk-withdegree, adds the node degree as an extra feature to the embeddings. Similarly, for the high-bias algorithm L3 [6], which counts paths of length 3 and applies a soft normalization, we create a lower-biased version called L3n by applying a stronger normalization based on node degrees. Besides these, we consider preferential attachment (a purely-biased baseline that specifically brings forward interactions between high degree nodes), LINE [36] a neural-network based embedding algorithm with high bias (since its learning process captures node degree information in its embeddings; S. Figure 2), two network propagation algorithms von Neumann [37] and random walks with restarts (RWR) [38], both having low-bias due to strong normalization based on node degrees in their formulation.

### View #1: Bias of link prediction algorithms toward high-degree proteins

- **Proposed strategy:** To understand the disposition of an algorithm toward well-studied proteins, measure its similarity with preferential attachment (biased baseline).
- Provides information about how much an algorithm prioritizes high-degree nodes. Node degree is considered an indicator of how well-studied a protein is.

Here, we aim to investigate and quantify the bias in the predictions of the algorithms toward well-studied proteins. For this purpose, we utilize preferential attachment as a biased baseline and quantify the similarity between the predictions of the algorithms and those of preferential attachment. We quantify this computing the overlaps for varying number of predictions (Figure 1b). By analyzing the area under the curves generated from this analysis, we obtain normalized scores where +1/0/-1 indicates bias towards high degrees/no bias/anti-bias towards low degrees. The results of this analysis are aligned with our expectations (Figure 1c): Common neighbors and L3 exhibit the highest bias, followed by Deepwalk-withdegree and Line. Other algorithms exhibit relatively lower bias, while Jaccard index and Deepwalk are slightly biased toward low-degree nodes.

### Standard settings for evaluating link prediction algorithms

Here, our goal is to investigate the impact of standard evaluation settings on bias in predictions towards well-studied entities in the context of network biology and proteomics [30]. For this purpose, following the recommendations of the machine learning community [18, 24], we employ two approaches for generating training and test sets: (i) Edge-Uniform: We sample 10% of the edges in the network (using 2020 version) uniformly at random and split them to form the test set, (ii) Across-Time: We use a more recent, 2022 version of the network as the test set while the older 2020 version serves as the training set.

In Figure 2, the precision-recall curve is shown for all algorithms for both benchmarking datasets. Additionally, we compute two metrics: The area under precision-recall curve (AUPR) and the area under precision-recall curve in log-log scale (AUlogPR). Link prediction problems involve a large background set, with the number of possible node pairs scaling quadratically with the number of nodes. As a result, even achieving a 10% recall rate corresponds to a substantial number of predictions (approximately 10^5^ to 10^6^ for edge-uniform and across-time data; S. Figure 3). Thus, AUPR in linear scale primarily measures the “late curve predictivity” as more than 90% of its effective region consists of a high number of predictions (*>* 10^5^). On the other hand, AUlogPR emphasizes lower recall values in logarithmic intervals, representing a lower number of predictions and serving as a measure of “early curve predictivity.” While recent literature employs metrics like early precision to assess early curve predictivity [39], AUlogPR offers an advantage as it does not require a fixed threshold to define the “early” region.

**Figure 2:**
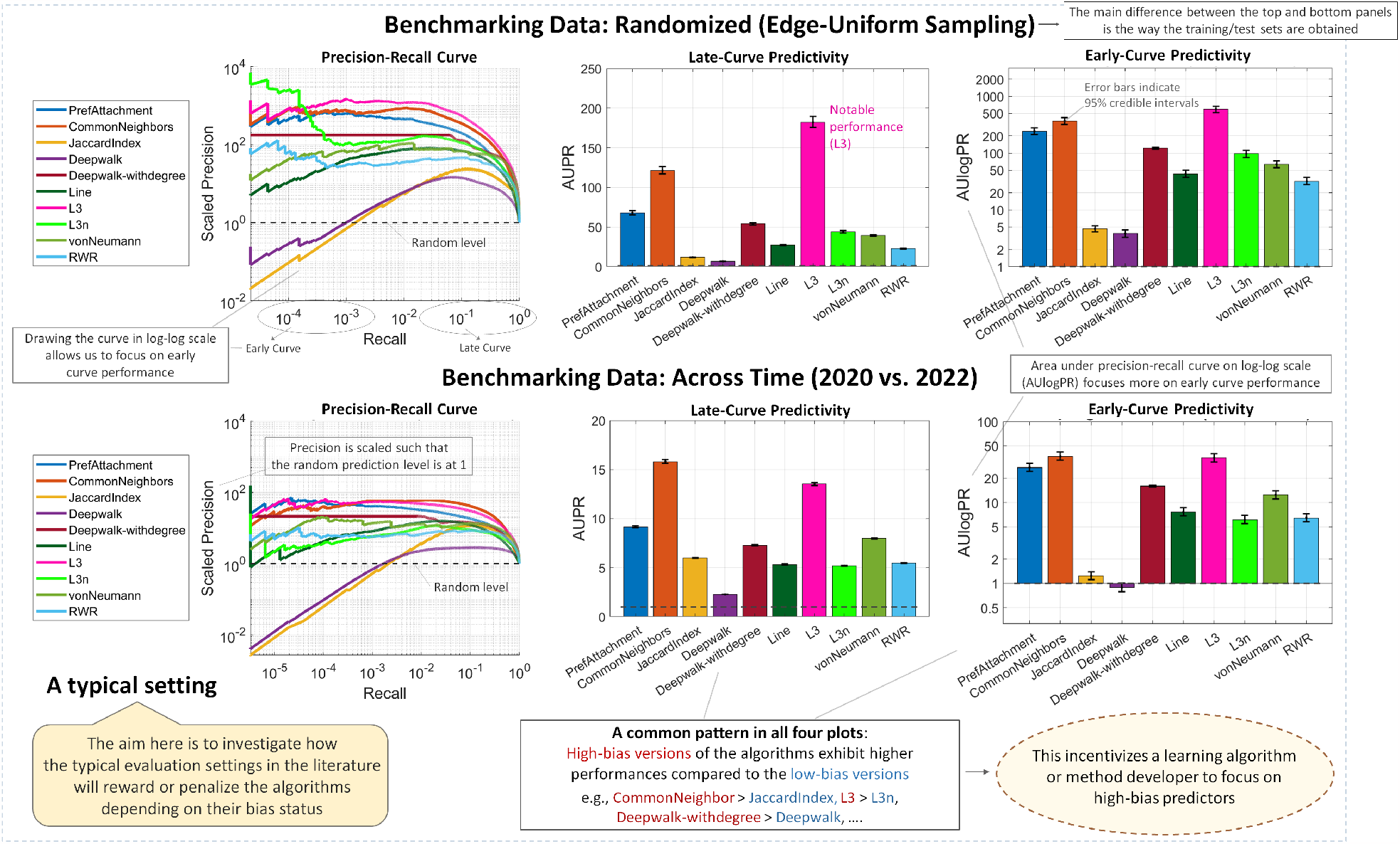
Results of a typical evaluation setting investigating the prediction performance of the link prediction algorithms in the context of PPI predictions. Two types of benchmarking data is considered: (Top panel) Randomized edge-uniform sampling and (Bottom panel) different snapshots of the network across time are used to generate the training and test instances.

As seen in Figure 2, the algorithms that are biased toward high-degree nodes seem to outperform other algorithms according to this evaluation setting, where the high-bias versions of the algorithms (CommonNeighbors, L3, Deepwalk-withdegree) exhibit considerably higher prediction performance compared to their low-bias versions (JaccardIndex, L3n, Deepwalk). Furthermore, the differences based on the degree bias appear to be more pronounced in the early curve (AUlogPR). Overall, these results show that the standard evaluation settings for evaluating link prediction algorithms can incentivize an algorithm or method developer to focus on high-bias predictors that bring forward well-studied biological entities at the expense of under-studied ones.

### View #2: Bias in benchmarking data and evaluation framework

- **Aim:** To assess bias in an evaluation setting by examining the incentive it provides to high-bias predictors favoring well-studied entities.
- **Proposed strategy:** Quantify the predictive power of node degrees in distinguishing positives from negatives in the test set, using preferential attachment as a representative model.

To understand the reasons behind the observed bias favoring high-degree nodes in the standard evaluation setting, we conducted an analysis of imbalance in the network and benchmarking data. In this analysis that we refer as separability analysis, we focus on the degree distribution and its predictive power in separating the “positives” (known interactions hidden from the algorithms) from the “negatives” (possible node pairs without a known interaction) with the help of Kolmogorov-Smirnov (K-S) statistic. In this analysis, we first categorized the nodes into three groups based on their connectivity, taking into account the cumulative degree distribution (Figure 3a): Poor nodes (≤ 20 interactions), Moderate nodes (20-100 interactions), and Rich nodes (*>* 100 interactions).

**Figure 3:**
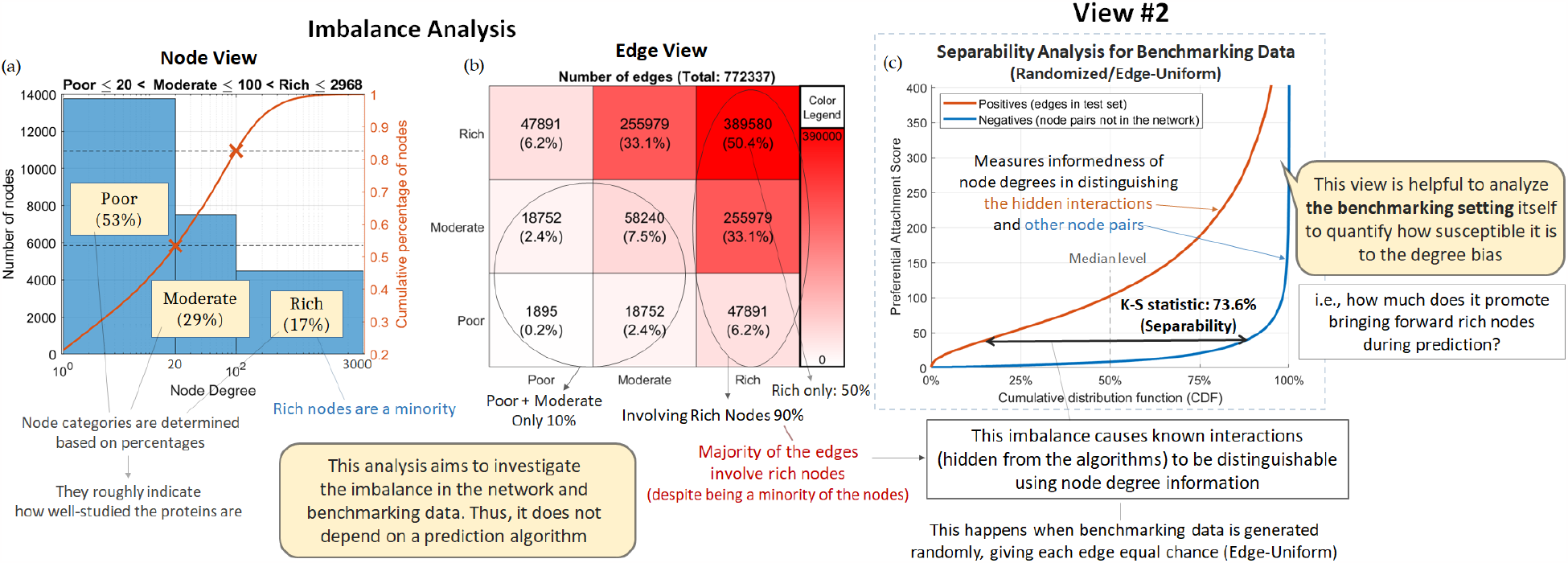
Investigating the imbalance in the benchmarking data and the incentive towards high-bias predictors by quantifying the predictive power of node degree information in distinguishing the known interactions on Biogrid PPI network. (a) Assigned node categories indicating how well-studied a protein is based on node degree information. (b) The distribution of the edges in the network across these categories. (c) Separability analysis for the randomized/edge-uniform setting.

After categorizing the nodes into three groups, we categorize the interactions in the network into 3x3 groups, encompassing all possible combinations of incident node categories (Figure 3b). This analysis reveals a significant imbalance in the distribution of edges across different node groups: Despite Poor and Moderate nodes constituting approximately 85% of all nodes in the network, 50% of all edges are between two Rich nodes and 90% of all edges involve at least one Rich node.

This substantial imbalance has important implications in the evaluation metrics. In standard settings, where all edges are equally valued and the edges in the test set are randomly sampled with uniform probabilities (Edge-Uniform), the evaluation heavily favors high-degree nodes. In fact, Rich nodes receive approximately 70% of the expected influence, while Poor nodes, despite comprising a majority (53%) of all nodes, receive only 5% influence. This discrepancy effectively incentivizes algorithms to prioritize the prediction of new interactions for well-studied proteins at the expense of under-studied ones. However, uncovering new interactions between under-studied proteins could potentially be more significant for generating novel insights in biological knowledge [31].

Figure 3c shows that the edges in positive set generated through random (edge-uniform) sampling are largely distinguishable from negative pairs using node degrees (K-S statistic: 73.6%). Even when multiple snapshots of the network across time are used for evaluation, the positive edges remain significantly separable from negative pairs (K-S statistic: 59.3%; S. Figure 4), This suggests that using a different snapshot of the network across time as a test set, as opposed to a randomly sampled test set, is not sufficient alone in addressing the issue of favoring biased predictions. Instead, it reinforces the finding that research tends to generate knowledge predominantly involving well-studied proteins [31], as nodes with high degrees in the earlier network tend to gain interactions over time.

Based on these observations, we conclude that an alternative evaluation approach is necessary to address the under-representation of under-studied proteins in evaluations. To tackle this issue, in the subsequent view, we focus on a simple idea: valuing each node equally, rather than each edge.

### View #3: Weighted evaluation setting focusing on under-studied entities

- **Aim:** To ensure that under-studied proteins are not under-represented while assessing the prediction performance of link prediction algorithms.
- **Proposed strategy:** Employ weighted metrics incorporating node degrees to assign greater importance and value to the discovery of interactions involving under-represented nodes.

Motivated by our previous observations, we propose a weighted evaluation setting to address the evaluation gap for under-studied proteins. This setting aims to balance the influence of each node in the evaluation process, resulting in a node-uniform evaluation. To achieve this, we formulate an optimization problem to minimize the difference between the weighted node degrees and a desired distribution. We develop an iterative algorithm (Algorithm 1) to solve this problem and optimize the weights assigned to each edge for uniform distribution. These optimized weights are then used as instance weights during the computation of evaluation metrics. Alternatively, these weights can be used as probabilities for generating node-uniform sampled test sets during evaluation.

The weighted metrics successfully balance the influence of nodes in the evaluation process, with an expected influence of 40%/35%/25% for Poor/Moderate/Rich nodes in the weighted setting (S. Figure 5). Additionally, they mitigate the degree bias in the benchmarking data by reducing the predictive power of node degree information, as indicated by the K-S statistics of 33.9%/18.4% for edge-uniform/across-time in the weighted setting (S. Figure 5d).

Figure 4 shows evaluation results of link prediction algorithms for across-time setting using the weighted metrics (results for sampled data can be found in S. Figure 6). These results differ significantly from those suggested by the standard settings shown in Figure 2. In this weighted setting, low-bias versions of the algorithms tend to outperform their high-bias counterparts. In addition, biased algorithms and excessively anti-biased algorithms, such as the anti-preferential attachment model, do not perform well in this setting (S. Figure 7). This indicates that the weighting approach effectively mitigates the degree bias without introducing an anti-bias effect by excessively inflating the weights of low-degree nodes.

**Figure 4:**
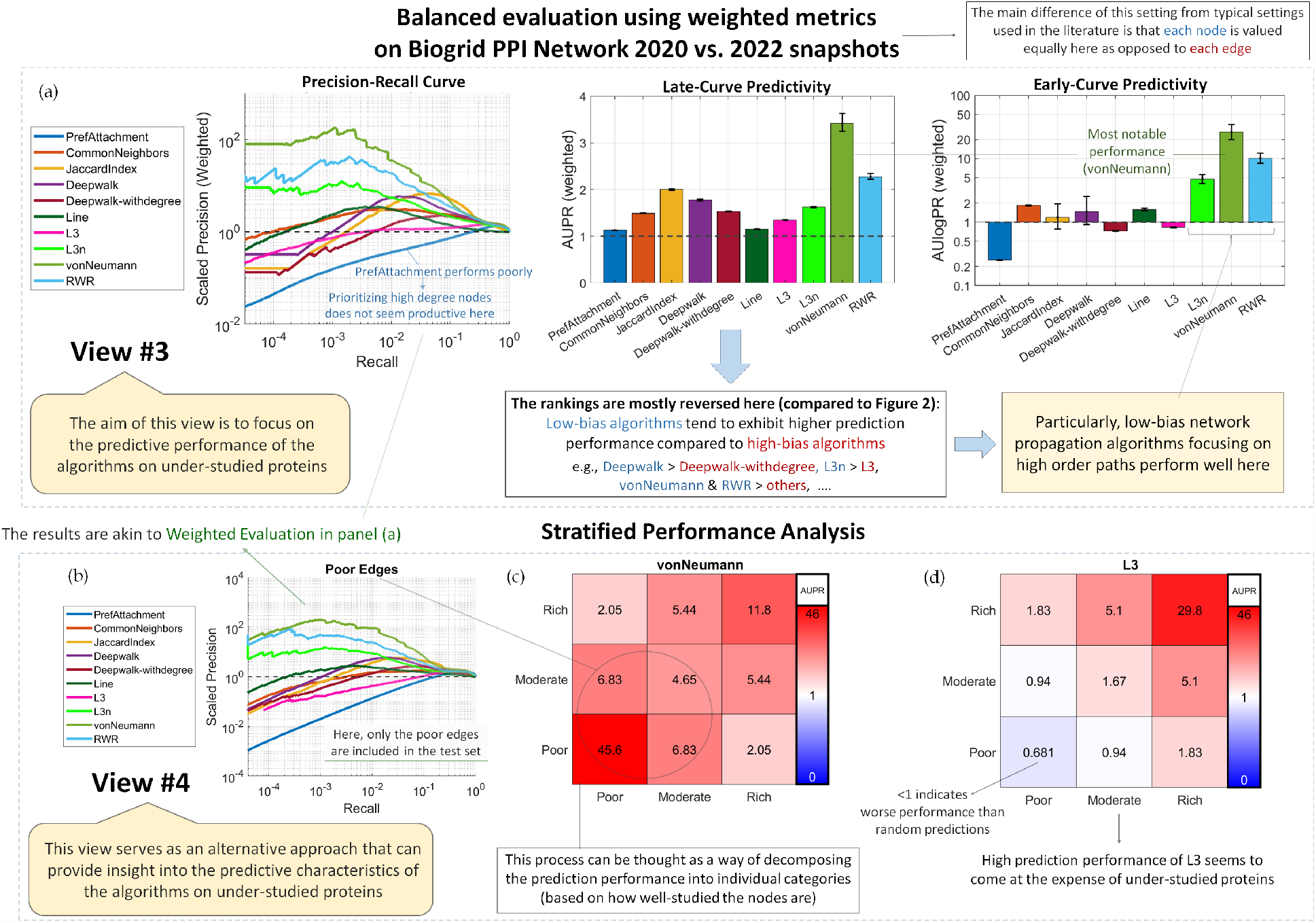
Balanced evaluation setting focusing on the prediction performance of the algorithms on under-studied proteins with the use of weighted metrics (node-uniform) and stratified analysis on Biogrid PPI Across-Time (2020 vs. 2022) data. Prediction performance of the algorithms for (a) weighted analysis and (b) stratified analysis. (c, d) Late curve predictivity (AUPR) stratified by node categories for von Neumann and L3 algorithms.

### View #4: Stratified analysis to focus on under-studied proteins

- **Aim:** To assess the prediction performance of the algorithms for uncovering new interactions depending on how well-studied the involved proteins are.
- **Proposed strategy:** Stratify prediction performance into edge categories based on the degrees of incident nodes, evaluating each category separately by including only the corresponding interactions in the test set.

To assess the prediction performance of the algorithms for uncovering new interactions involving under-studied proteins in a more direct manner, we propose a stratified approach based on node degrees. This approach involves categorizing the edges into different categories based on the node degrees and assessing the prediction performance within each category. We stratify the edges into 3x3 categories, as depicted in Figure 3b, and further group them into two categories: Poor edges (edges between Poor+Moderate nodes), and Rich edges (remaining edges involving a rich node).

In the analysis of across-time data, the precision-recall curves for Poor edges, as shown in Figure 4b, are quite similar to those obtained through weighted analysis (Figure 4), but distinct from the standard setting (Figure 2). This is expected since Poor edges are assigned 54% influence in the weighted setting compared to 8% in the unweighted setting (S. Figure 8). Similarly, the curves for Rich edges closely resemble those in the standard setting, as these edges are given 92% influence there (S. Figure 9). To further examine the performance of the best performing algorithms in the stratified analysis, we present the results for vonNeumann and L3 in Figure 4c-d. Notably, vonNeumann exhibits relatively consistent predictivity across different edge categories. In contrast, while L3 demonstrates high predictive performance for Rich-Rich and Rich-Moderate interactions, it appears to come at the expense of severely diminished predictivity for edges involving understudied proteins.

### View #5: Diverse metrics for bias-aware comparisons

- **Aim:** To make a comprehensive and bias-aware evaluation in a simple manner.
- **Proposed strategy:** Measure five aspects, early/late curve prediction performance for under-studied/well-studied nodes, and the disposition of an algorithm regarding degree bias.

To facilitate bias-aware evaluation while striking a reasonable balance between simplicity and comprehensiveness, we propose five key metrics to summarize and compare the prediction characteristics of the algorithms (see Figure 5). These metrics assess the fairness of the predictions (defined as 1 minus absolute value of the bias metric), performance on well-studied entities (results of the standard/unweighted setting, valuing each edge equally), based on early and late curve characteristics measured by AUPR and AUlogPR, and the same metrics for under-studied entities (results of the weighted setting, valuing each node equally).

**Figure 5:**
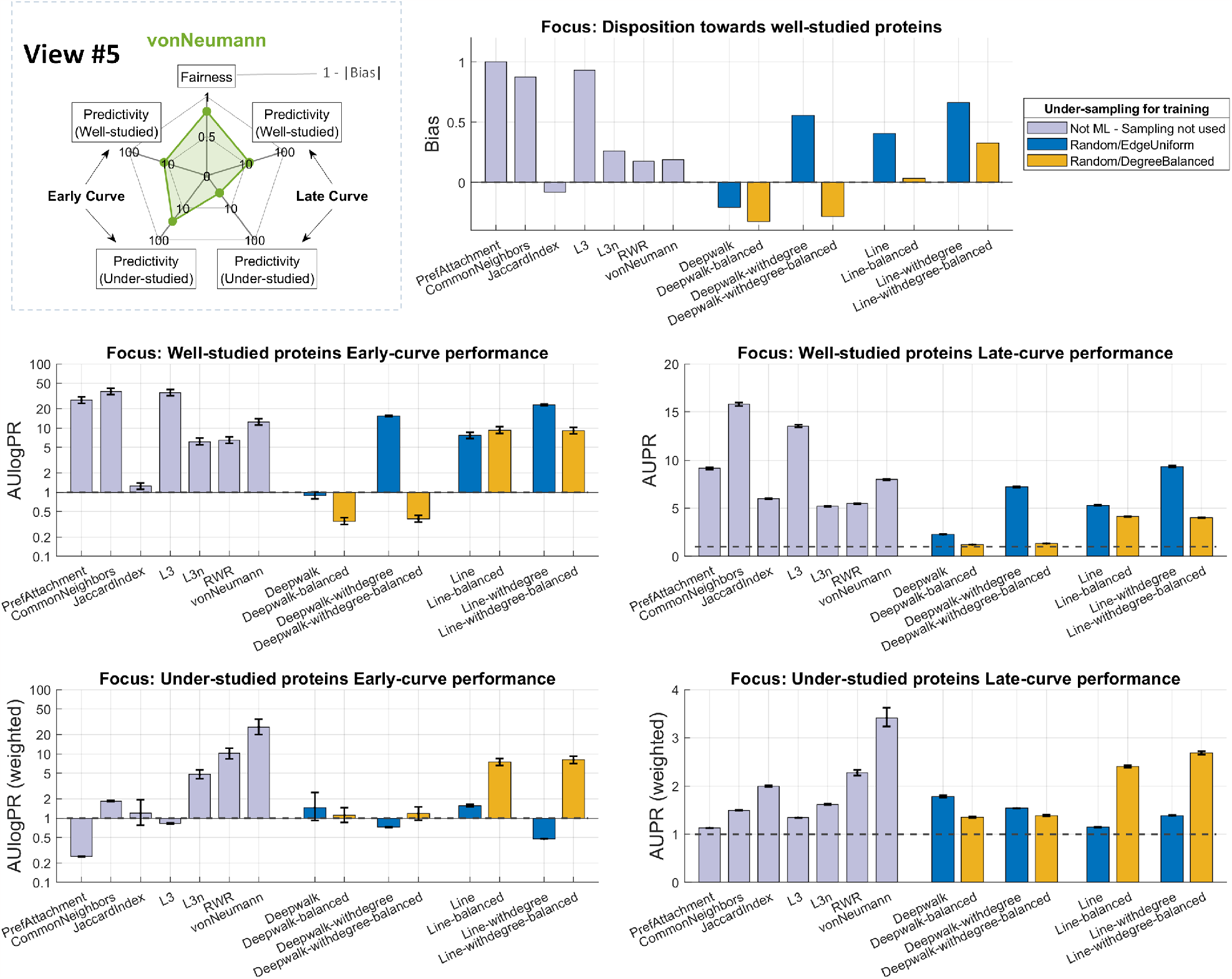
Results of bias-aware analysis on Biogrid PPI Across-Time (2020 vs. 2022). (Top Left) 5-metric summary for von Neumann algorithm. (Remaining Panels) Evaluation results for each of the five metrics are shown with bar plots: Bias (based on similarity with preferential attachment), AUPR, AUlogPR, weighted AUPR and weighted AUlogPR to focus on under-studied proteins. In each panel, the coloring is done according to the type of the sampling strategy used to cope with imbalance during the training of the algorithms.

### Exploring bias-aware training: Degree-balanced under-sampling strategy

To address bias in link prediction algorithms, we explore a Random/Degree-Balanced undersampling strategy during training. This approach aims to achieve balanced degree distributions for positive and negative samples and mitigate degree imbalance.

In our experiments, we compare the performance of DeepWalk and LINE, along with their biased variants incorporating node degrees (DeepWalk-withdegree and LINE-withdegree). Using the Biogrid PPI dataset in the Across-Time setting for 2020 vs. 2022, our results in Figure 5 show that the degree-balanced approach is effective for reducing bias in predictions towards well-studied proteins. Additionally, we observe a significant improvement in the prediction performance of LINE for understudied proteins, as measured by the introduced weighted metrics. In contrast, standard evaluation metrics focusing on well-studied proteins indicate decreased performance, underscoring the need for explicit evaluation and measurement of performance on understudied proteins.

Note that, the impact of degree-balanced sampling on the performance of understudied proteins appears to depend on whether it results in an anti-bias effect. When combined with biased algorithms like LINE, the degree-balanced sampling strategy results in reduced bias and improved performance for understudied proteins. On the other hand, when applied to already low-bias algorithms such as DeepWalk, it leads to an anti-bias effect and does not translate into improved performance. This observation highlights the importance of assessing algorithm bias in determining the effectiveness of specific sampling strategies.

### Comprehensive evaluation on different types of PPI networks and settings

To comprehensively analyze biases in different types of PPI networks, we conducted additional analyses using Biogrid datasets from multiple time points (2006, 2010, 2015) versus the 2022 dataset in an Across-Time setting (S. Figures 10-12). Additionally, we explored various evidence subnetworks within the STRING database [40], including experimental, text mining, coexpression, and the combined network. We performed experiments in both the Random/Edge-Uniform setting (S. Figures 13-16), Across-Time setting (2015 vs. 2021; S. Figures 17-20), as well as an Across-Evidence setting (experimental vs. combined; S. Figure 21).

Our findings reveal a progressive increase in imbalance across different time settings in older Biogrid networks. The K-S statistic indicates an increase from 33% in the 2006 vs. 2022 setting to 59% in the 2020 vs. 2022 setting. Among the tested network types, random/edge-uniform exhibited the highest levels of imbalance, with K-S statistics ranging from 50-80%, while the across-evidence strategy in STRING showed the lowest imbalance with a K-S statistic of 30%. We also observe that weighted evaluation effectively addressed the bias, resulting in a reduction in K-S statistic by up to 50%, to values typically less than 15%. Overall, these results are aligned with our previous conclusions, further reinforcing the significance of our observations.

### Bias in benchmarking data for other link prediction problems in biology

To assess the generalizability of our findings to other link prediction problems in biomedical applications, we investigate the bias in benchmarking data for several prominent networks (Figure 6). Specifically, we investigate PhosphoSitePlus kinase-substrate interactions (PSP-KS) [41], TRRUST transcription factor regulatory interactions [42], DrugBank drug-drug interactions (Drugbank-DDI) [43], and drug-disease association (NDRFT-DDA [1] and CDA-DDA [44]) networks.

**Figure 6:**
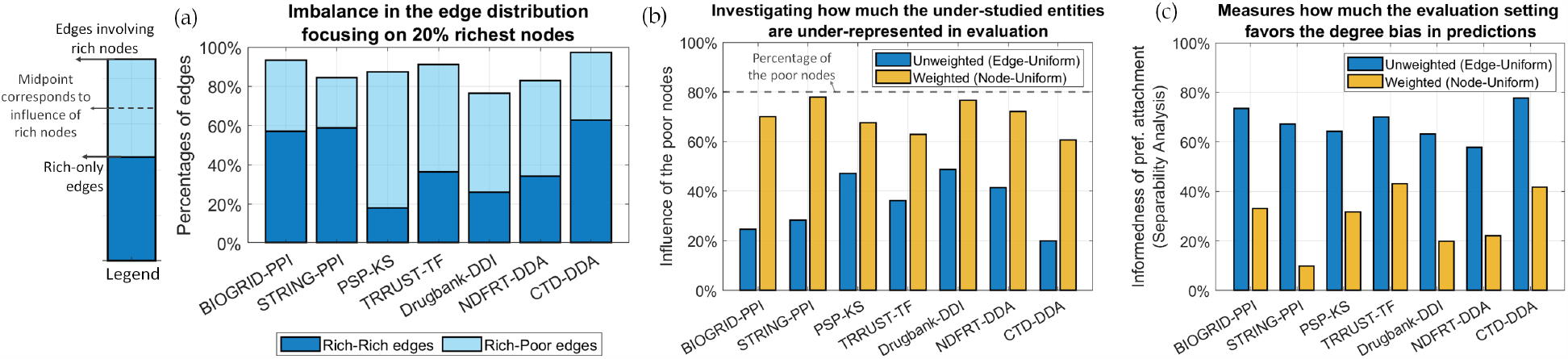
Investigating different datasets and link prediction problems in the context of network biology in terms of the imbalance in the network, under-representation of the understudied entities in the evaluation, and the incentive towards high-bias predictors. (a) The percentage of edges involved with the 20% richest nodes in the network. (b) Influence of the under-studied (i.e., 80% lowest degree) nodes in the evaluation for weighted (node-uniform) and unweighted (edge-uniform) settings. (c) Predictive power of the node degree information. Visit https://yilmazs.shinyapps.io/colipe to investigate the bias in these datasets interactively.

For each network, we examine the imbalance in the edge distribution for the top 20% of the nodes with highest degrees (Figure 6a). Next, we consider both the weighted (node-uniform) and unweighted (standard, edge-uniform) settings and quantify the under-representation of understudied entities in the evaluation by measuring the influence of the 80% of nodes with the lowest degrees (Figure 6b). Additionally, we assess the incentive provided by the evaluation setting towards high-bias predictors by measuring the predictive power of node degree information Figure 6c). Our analysis reveals that a wide range of network datasets commonly used for benchmarking link prediction algorithms exhibit a significant degree of imbalance in the edge distributions. As a result, standard evaluation settings that assign equal value to each edge tend to reward algorithms that prioritize high-degree, well-studied entities in their predictions.

## Discussion

Our brief, albeit non-exhaustive, exploration of recent literature in biomedical context suggests there are more than 109 link prediction papers published within the last five years alone (published 2018-2023; a list of these papers is provided in the supplementary materials). The applications in these papers encompass a diverse array of problems, including the problems discussed in the final part of our analysis (Figure 6). This high volume of research is a testament on the growing significance of machine learning in LP tasks within the field of biomedicine, and this trend is bound to gain further momentum in the upcoming years.

Our study uniquely contributes by addressing a critical gap: the incorporation of bias-aware evaluation, an often overlooked facet in this fast evolving field. Our results across various biomedical applications and network datasets reveals a notable imbalance in edge distributions, favoring high-degree, well-studied entities in standard evaluation settings. These findings underscore the need for bias-aware evaluation strategies to achieve a comprehensive assessment of algorithm performance and facilitate the discovery of interactions involving under-studied entities. While our analysis centers mainly on PPI predictions, the developed techniques are applicable beyond the protein domain.

Overall, our study and the strategies we present aim to address the issues arising from the severe imbalance in standard evaluation settings within biological networks. To promote bias-aware evaluation and enhance exploration of new interactions in these networks, we propose the following approaches for developers and reviewers of link prediction algorithms, organized in an easy to remember acronym AWARE:

- **Analyze bias in algorithms:** To uncover algorithms’ disposition towards well-studied nodes, investigate their similarity with predictions of preferential attachment model (View 1).
- **Watch out for bias in benchmarking data:** Examine the benchmarking setting to quantify potential imbalances and the extent to which it rewards high-bias predictors (View 2). Explore alternative evaluation settings, such as Across-Evidence and Across-Time, for a more realistic assessment, going beyond random sampling.
- **Assess prediction performance on under-studied entities:** Adopt a weighted setting, assigning value to each node as opposed to each edge equally (View 3), or conduct a stratified analysis (View 4) to evaluate the prediction performance on under-studied proteins.
- **Review diverse metrics to draw conclusions:** Consider several aspects to outline the main characteristics of an algorithm, encompassing early curve/late curve performances, well-studied/under-proteins, as well as the bias in predictions (View 5).
- **Engage in bias-aware training:** Recognize the impact of an algorithm’s inclination towards well-studied entities on its predictions. To develop innovative algorithms prioritizing understudied entities, incorporate effective normalization techniques or employ degree-balanced sampling strategies during the training of link prediction algorithms to address the biases inherent in the data.

We anticipate that the adoption of these AWARE principles will aid the research community in gaining better insights from computational evaluations and promote the development of targeted algorithms for under-studied proteins, facilitating new discoveries.

## Materials and Methods

First, we provide an overview of the methodology we propose without the implementation details, focusing on evaluation metrics, the proposed weighted validation setting, and the degree-balance sampling strategy. Technical descriptions and other information regarding the experimental details are provided in their appropriate sections below.

### Evaluation metrics

To measure the late-curve prediction performance, we utilize the area under the precision-recall curve (AUPR). For the early-curve performance, we compute the area under the precision-recall curve in log-log scale (AUlogPR) through numerical integration (after normalizing the logarithmic x-axis such that the resulting unit for AUlogPR is precision). Note that, to make the evaluation results comparable with different networks or settings, we scale both metrics to have an expected value of 1 for random predictions. To account for the variance in the estimation of these meaures, we construct 95% credible intervals following a Bayesian approach [45].

### Optimization algorithm for node-uniform edge weights

To obtain a set of edge weights (denoted as **W** matrix) that establishes node-wise uniformity (i.e., for the row and column sums of **W** to be equal for all nodes), we formulate this as an optimization problem and develop an algorithm that iteratively performs multiplicative updates (ensuring the uniformity of the rows in one step, and for the columns in another) until the uniformity of the row and column sums are established simultaneously at an acceptable level. The formulation of the optimization problem and a simplified pseudo-code of the developed algorithm (omitting some details) is given in Algorithm 1. A more detailed description of the algorithm (including a complete pseudo-code) specifying the technical details (e.g., regarding matrix initialization, termination conditions, controlling the step size during updates and so on) are provided in the *“Weighted Validation Setting”* section and Algorithm 3. Note that, we denote **Q**_**r**_ and **Q**_**c**_ to indicate the desired weights for rows and columns instead of assuming uniformity for the sake of generalizability (i.e., node-uniform when **Q**_*r*_ = **Q**_*c*_ = **1**).

#### Algorithm 1 Optimization Problem and Algorithm (Simplified Pseudo-Code)

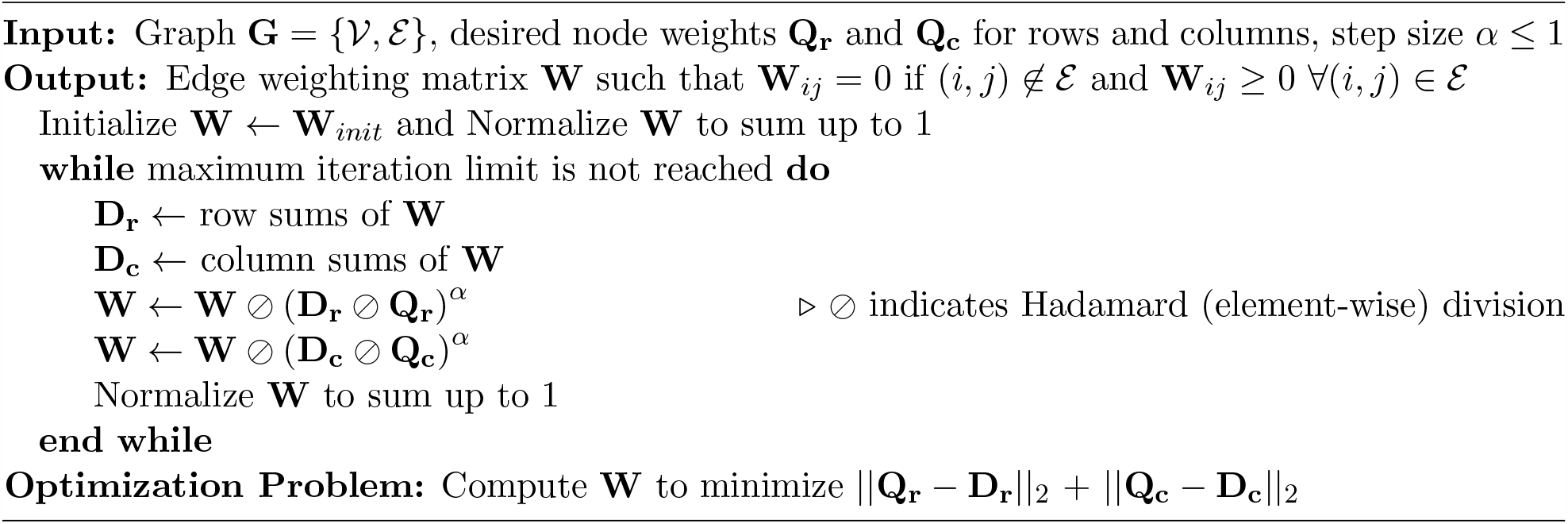

### Weighted evaluation

In the weighted evaluation setting, we use the optimized edge weights (the matrix **W** computed by the algorithm above) as the weight for each positive instance (existing edge in the test set). For this purpose, we generalize the computation of AUPR and AUlogPR to assign weights to instances (positives in the test set) while counting the number of true positives (TPs) and false negatives (FNs). For example, an edge in the test set that is weighted worth of 3 unweighted edges, if included in the predictions of an algorithm, would increase the number of TPs by 3 as opposed to 1. Performance measures are then computed based on these weighted counts.

### Degree-balanced under-sampling strategy for training

In this section, we aim to create a balanced training set for machine learning algorithms by ensuring that the size and node degree distribution of the under-sampled negative set (node pairs) match those of the positive set (edges in the training network). To achieve this, we adopt a random sampling approach where the selection probabilities of node pairs are assigned based on the degrees of the nodes in the network. For more details on the methodology, please refer to Algorithm 2. Note that, in this context, we assume the network to be undirected, meaning that adding a node pair (*i, j*) is equivalent to adding (*j, i*) to the network.

#### Algorithm 2 Random/Degree-Balanced Under-Sampling Algorithm

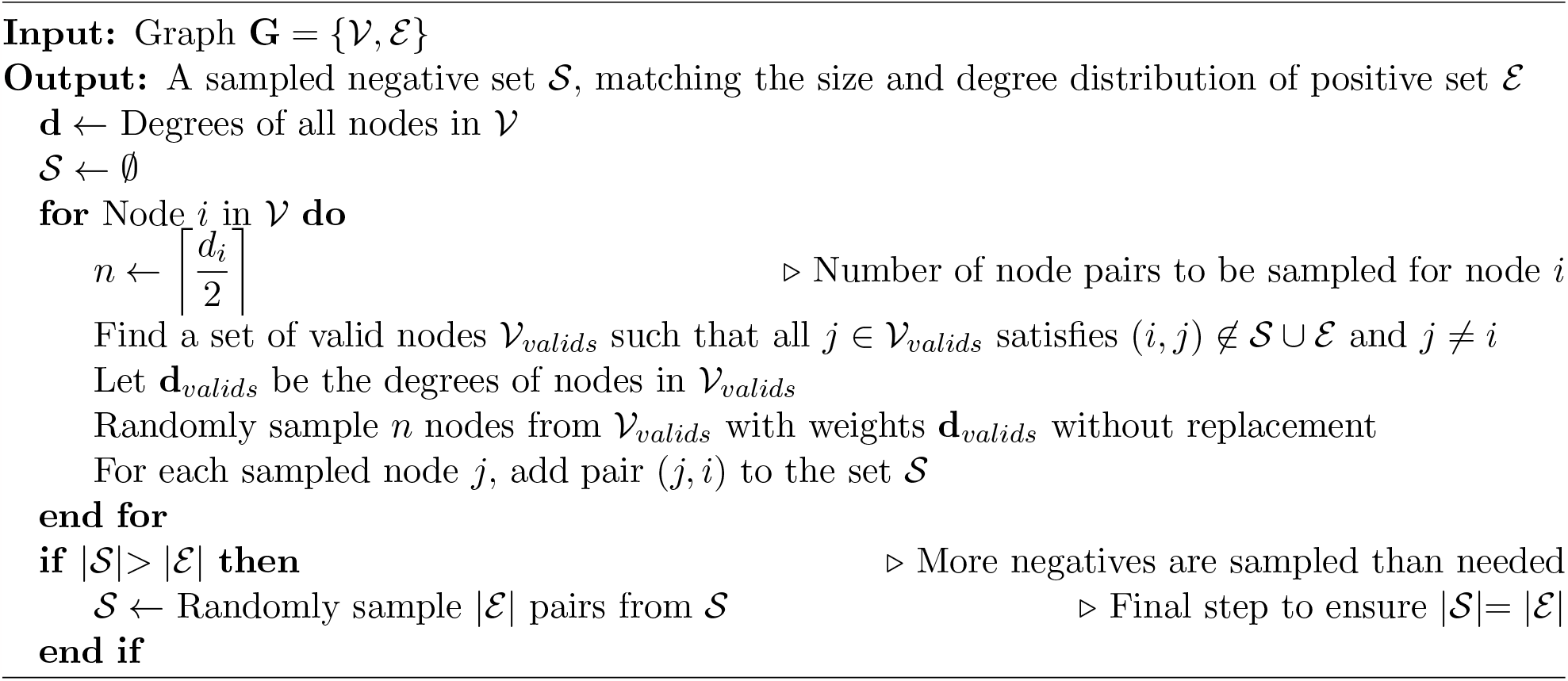

### Computing the influence of the nodes on evaluation

We quantify the influence of node categories (Poor/Moderate/Rich) on the evaluation by examining the percentages of edges belonging to each category, as illustrated in Figure 3b and S. Figure 5b. The influence is determined based on the number of edges in the standard unweighted setting, or the weights of edges in the weighted evaluation setting. In the case of mixed category edges, their influence is divided equally between the respective categories. For example, a Poor-Rich edge contributes half of its weight to the Poor category and the other half to the Rich category. This approach ensures that the total influence of the node categories adds up to 1.

### Quantifying the bias towards high-degree nodes in method predictions

Here, we assess the overlap between the predictions of each algorithm and the preferential attachment model by counting the number of shared positive predictions for different values of *k*, where *k* represents the number of top predictions for both algorithms. The bias metric is then computed based on the area under these curves in log-log scale and normalized to fall within the [-1, 1] range. This normalization is done according to the maximum value corresponding to *k*, and the expected value for random predictions, which is calculated as *k*^2^ divided by the total number of pairs to be predicted.

### Separability analysis quantifying the degree bias in benchmarking data

In this analysis, we compute the preferential attachment scores based on the node degrees in the training data for both the positive interactions (hidden interactions in the test set) and negative node pairs (pairs without known interactions). We quantify the predictive power of node degree information by using the Kolmogorov-Smirnov statistic, which measures the maximum distance between the cumulative distribution functions. This approach is equivalent to computing the informedness [46] of the preferential attachment model at its best prediction point, where the receiver operating characteristics (ROC) curve has the maximum vertical difference from the diagonal line representing random predictions. For the weighted version of the separability analysis, we use the optimized edge weights as instance weights while estimating the cumulative distribution function (CDF) for the positive interactions and compute the Kolmogorov-Smirnov statistic using the CDF for the weighted positives.

### Performing stratified analysis

For the stratified analysis, we first obtain the node categories (Poor, Moderate, Rich) as shown in Figure 3a. We assign these categories by considering the cumulative degree distribution so that Poor and Rich nodes roughly comprise 50% and 20% of all nodes in the network. Next, we assign each edge in the test set into one of six categories (e.g., Rich-Rich, Poor-Rich, Poor-Moderate and so on). For each of these categories separately, we repeat the evaluation (and compute performance metrics like AUPR), keeping only the edges in the corresponding category in the test set (in other words, considering the prediction of only the edges in that category to be true positives). Note that, the background set of possible node pairs is not affected by this stratification i.e., the negative set includes pairs from all categories.

### Datasets used in this work

The bulk of the experiments done in this paper uses BioGRID [35] Human Protein-Protein Interaction network for two versions obtained at different times: (i) 2020 version (v4.0.189) contains 464,003 interactions between 25,776 proteins, (ii) 2022 version (v4.4.210) contains 784,774 interactions between 27,408 proteins. For constructing the training/test sets across time (2020 for training, new edges in 2022 for testing), we filter for the proteins that exist in the 2020 version and use the 308,334 new interactions for 16305 proteins in 2022 version as the test set (for a total of 772,337 interactions between 25,776 proteins, training & test sets combined).

The final part of our analysis includes six other networks listed below. Some of them were obtained and parsed from the source databases directly, while others are taken from BioNEV [1] repository as pre-processed edgelists.

- STRING PPI [40] contains 359,776 interaction between 15,131 proteins. Taken from BioNEV repository as an unweighted undirected network.
- PhosphoSitePlus Kinase-Substrate (PSP-KS) dataset [41] contains 13,664 Kinase-phosphosite pairings. Taken from source (PhosphoSitePlus) and parsed by us as an unweighted undirected heterogeneous bipartite network. Filtered only to contain pairs observed in Human tissue.
- TRRUST [42] (Transcriptional Regulatory Relationships Unraveled by Sentence based Text mining) dataset contains 3,149 transcription-factor relationships between 1,621 genes. Taken from source and parsed by us as an unweighted undirected network. Filtered only to contain Activation relationships (as opposed to repression or unknown).
- Drugbank [43] Drug-Drug Interaction dataset contains 242,027 interactions between 2,191 drugs. Taken from BioNEV repository as an unweighted undirected network.
- NDFRT is a Disease-drug association dataset containing 56,515 associations between 13,545 diseases and drugs. Taken from BioNEV repository as an unweighted undirected heterogeneous bipartite network.
- CTD [44] is a Disease-drug association dataset containing 92,813 associations between 12,765 diseases and drugs. Taken from BioNEV repository as a pre-processed unweighted undirected heterogeneous bipartite network.

### Biogrid human PPI across-time setting on older datasets 2006-2015

We also assess the effect bias in the across-time setting by examining older networks from Biogrid. Specifically, we analyze the Biogrid 2006 (v2.0.17) network, the earliest release accessible on the Biogrid website, which comprises 20,020 interactions involving 7,368 proteins. Additionally, we investigate the Biogrid 2010 network (v3.0.68) with 32,347 interactions among 9,715 proteins, and the Biogrid 2015 network (v3.3.124) encompassing 171,711 interactions among 20,749 proteins. In each case, we utilize the new interactions from the recent 2022 version (v4.4.210) as the test set.

### STRING PPI evidence subnetworks, across-time and across-evidence settings

Additionally, we explore various types of networks within the STRING human PPI network where the type of a network refers to the evidence used to identify the interactions in the network. These network types include the experimental, text mining, coexpression, and combined networks. We conduct experiments on each of these networks using two different settings: Random/Edge-Uniform and Across-Time. For the Random/Edge-Uniform setting, we utilize the latest release from 2021 (v11.5). It is important to note that STRING provides interaction confidence scores rather than the network itself. To construct the network, we apply a confidence threshold of 0.7, resulting in the following numbers of interactions and proteins for each subnetwork: 49,900 interactions among 8,012 proteins in the experimental subnetwork, 69,601 interactions among 15,339 proteins in the text mining subnetwork, 25,293 interactions among 2,418 proteins in the coexpression network, and 252,984 interactions among 16,814 proteins in the combined network.

For the Across-Time setting, we employ an older release from 2015 (v10) as the training set for the algorithms, and the latest release from 2021 (v11.5) as the test set for evaluation. In the older release from 2015 (v10), using a high confidence threshold of 0.7, we obtain the following numbers of interactions and proteins for each subnetwork: 39,551 interactions among 8,832 proteins in the experimental subnetwork, 65,429 interactions among 13,322 proteins in the text mining subnetwork, 16,846 interactions among 1,160 proteins in the coexpression network, and 321,152 interactions among 15,554 proteins in the combined network.

In addition, we explore an alternative setting called Across-Evidence, where we train the algorithms using the experimental subnetwork and evaluate their performance on the combined network. To conduct this analysis, we utilize the latest networks from 2021 (v11.5). This approach allows us to utilize 80,316 new interactions in the combined network as the test set, involving the 7,087 proteins that are present in both the experimental and combined subnetworks.

### Link Prediction Algorithms - Verbal Descriptions

#### Methods based on scoring metrics

We consider the preferential attachment model to represent a purely biased model (where node pairs are ranked based on the product of the degrees of the endpoints). Common neighbors represents a high-bias algorithm that considers paths of length 2 (high bias since the number of paths in correlated with the node degrees). Jaccard Index represents a low bias version of common neighbors where a normalization is applied based on node degrees.

#### High order paths/Network propagation algorithms

L3 is a method that counts the paths of length 3 to make predictions. For this purpose, in this work, we consider the formulation given in [6] that applies a soft normalization (based on square root of degrees, this is what we consider the high-bias version). We also introduce a low bias version of it, L3-Normalized (L3n) that applies a stronger normalization based on node degrees. Whereas, von Neumann [37] and random walks with restarts (RWR) [38] are network propagation algorithms that consider a weighted combination of paths of different lengths. Formulation of both algorithms involve a strong normalization based on node degrees (the main difference between them is the style of the normalization, whether it is done symmetrically or based on column normalization). Thus, we consider both to be low-bias algorithms.

#### Embedding/Learning Methods

We consider two types of embedding methods: Deepwalk [13] (Random walk based) and Line [36] (Neural network based). For each of these methods, we train a logistic regression model using the embeddings as features. Here, deepwalk represents a low-bias algorithm (since the embedding dimensions are uncorrelated with node degrees, likely by design, S. Figure 1) and Line is a higher bias algorithm (since its embeddings pick up the node degree info during learning, S. Figure 2). For deepwalk-withdegree, we include the node degrees as an additional dimension (as if it is part of the embeddings matrix) to construct a high-bias version of the deepwalk algorithm.

For each embedding algorithm, we train a logistic regression model to obtain predictions.While training the model, to ensure a balanced training set, we randomly sample the negative samples (i.e., node pairs that are not in the training set) to have the same size as the edges with positive labels. Unless otherwise specified, this under-sampling is performed as commonly applied in the literature [1, 13], using Edge-Uniform probability distribution, giving each node pair equal chance of being sampled. In addition to this, we also propose and explore with a new sampling approach that we refer as Random/DegreeBalanced. This approach performs random under-sampling of the negative examples while aiming to balance the degree distribution in both the positive and negative edge classes in an efficient manner. For further details and mathematical descriptions of the link prediction algorithms, please refer to the supplementary materials.

### Evaluation metrics - AUPR and AUlogPR

For computing the area under precision-recall curve (AUPR) and the log-log scale precision-recall curve (AUlogPR), we perform numerical integration as follows:

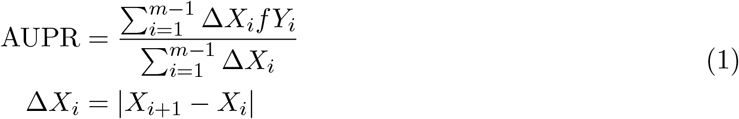

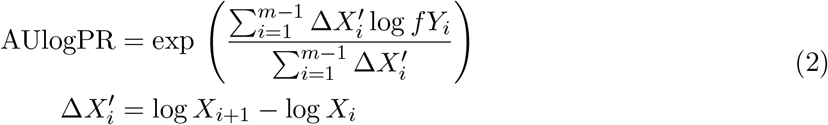

where *X*_*i*_ is recall and *Y*_*i*_ is precision values corresponding to the *i*th measurement point, Δ*X*_*i*_ is the gap between two consecutive points in the x-axis and *f Y* is an interpolating function for the area between two consecutive points:

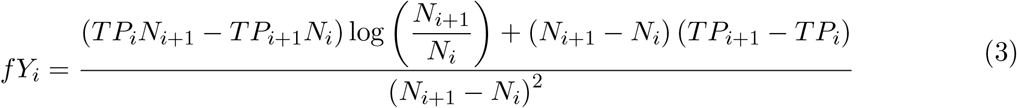

where *TP*_*i*_ denotes the number of true positives, *FP*_*i*_ the number of false positives, and *N*_*i*_ = *TP*_*i*_ + *FP*_*i*_ the number of predictions. Here, *f Y*_*i*_ function is an interpolating function used to calculate the normalized area between two consecutive points *Y*_*i*_ and *Y*_*i*+1_ on the precision-recall curve. It addresses the inaccuracy that can arise when there are large gaps between these points, especially for methods with discrete scoring. A simple linear interpolation, such as 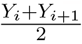, may not capture the precise relationship between precision and recall. To overcome this, we employ a customized interpolation function tailored for precision-recall curves. This interpolation helps reduce inaccuracies, particularly in cases with significant gaps between points and when working with discrete scoring or logarithmic scale. Further mathematical descriptions and derivations can be found in the supplementary materials.

Note that, while computing AUlogPR, we begin the curve at the *TP* = 10 point to mitigate the volatility observed in the initial points between *TP* = 1 and *TP* = 10. This step helps reduce variance in the estimation. After calculating the area under the curve for both AUPR and AUlogPR metrics, we normalize the values to have an expected value of 1 for random predictions. This normalization is achieved by dividing the calculated area by the prevalence of positive labels.

After computing the AUPR and AUlogPR metrics, we employ a Bayesian approach to construct credible intervals that reflect the variance in the estimation. For a detailed explanation on the construction of these intervals, please refer to the supplementary materials.

### Weighted Validation Setting

#### Optimization algorithm for obtaining edge weights based on node valuations

We formulate this problem as follows: Suppose we are given as set of node valuations *q*. Let *q*_*r*_(*u*) and *q*_*c*_(*u*) denote the desired expected number of edges coming into and going out of *u* ∈ 𝒱 (i.e., the desired row and column sums). Let **W** represent the weights of the edges as a sparse matrix where **W**_*ij*_ = 0 if (*i, j*) ∉ ℰ. Here, our aim is to estimate a set of edge weights/values **W** such that the row and column sums of **W** are respectively equal to *v*_*r*_ and *v*_*c*_. For this purpose, we will use an expectation-maximization based optimization algorithm with multiplicative steps. The pseudo-code of the algorithm is given in Algorithm 3. Below, we describe each step of the algorithm:

##### Initialization

In the beginning of the algorithm, we set **W** to be equal to an initial, approximate solution:

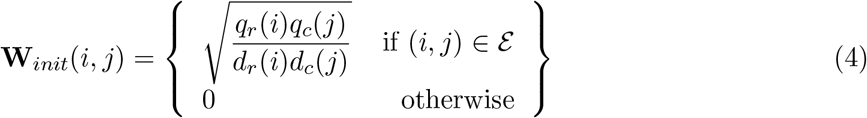

where *d*_*r*_(*i*) and *d*_*c*_(*j*) are the row and column degrees of nodes *i* and *j* in the network respectively. After setting **W** = **W**_*init*_, we normalize the weights **W** to sum up to 1 as follows:

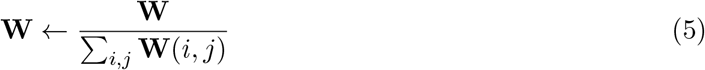

##### Update steps of the algorithm

Here, to ensure that the updated weights remain positive, we use multiplicative update steps based on row/column normalizations:

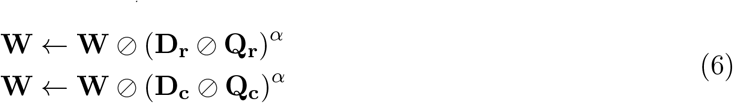

where *α* is a multiplicative step size parameter and **D**_**r**_ and **D**_**c**_ are row/column sum matrices of **W** respectively:

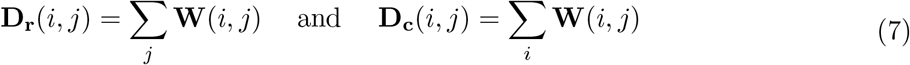

###### Algorithm 3 Optimization Algorithm for Edge Weights

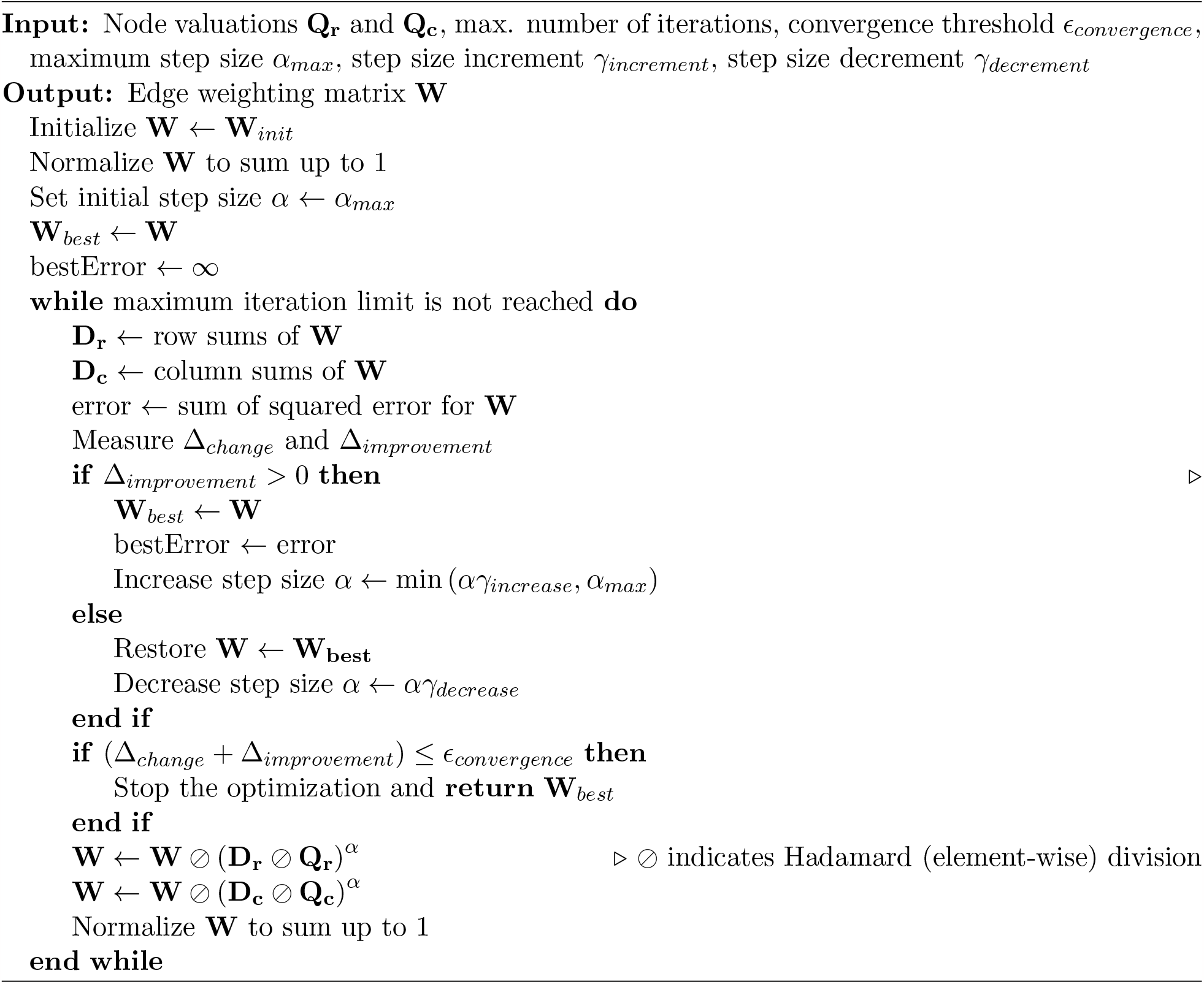

Similarly, input node valuation vectors *v*_*r*_ and *v*_*c*_ are organized as matrices **V**_**r**_ and **V**_**c**_ after being normalized:

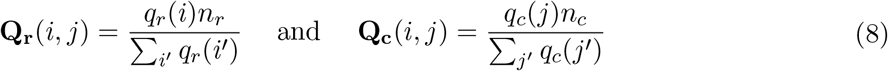

where *n*_*r*_ and *n*_*c*_ are scalars indicating the number of rows and columns in the network. In each step, after updating **W** according to Equation 6, **W** is normalized again to sum up to 1 as in Equation 5.

##### Termination of the algorithm

To determine the convergence of the algorithm, we look at two criteria. The first one focuses on the amount of change in **W**:

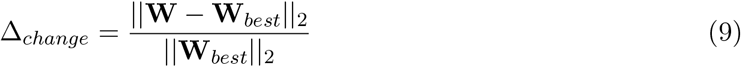

The second one focuses on the amount of improvement. For this purpose, we first quantify the error using sum of squares:

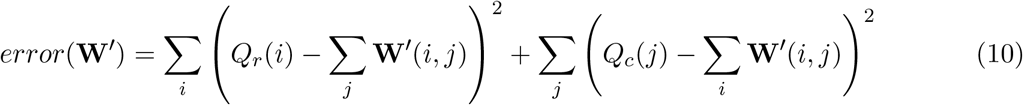

Thus, at each step, we measure the improvement **W** provides over **W**_*best*_ as follows:

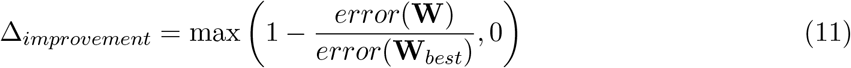

Overall, we terminate the algorithm when the amount of change plus the improvement is less than a predefined threshold *ϵ*_*convergence*_:

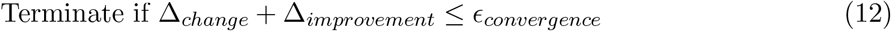

In addition to the convergence threshold, we terminate the optimization if the maximum iteration limit is reached. Unless otherwise specified, we use *ϵ* = 10^*−*2^ and 100 maximum iterations for the termination of the algorithm.

##### Updating the step sizes

When there is no improvement at any point (i.e., Δ_*improvement*_ ≤ 0), we conclude that step size is too large and need to be reduced. For this purpose, we restore **W** to **W**_*best*_ (i.e., the best weights with lowest error up to this point) and decrease step size *α* by a factor of *γ*_*decrease*_:

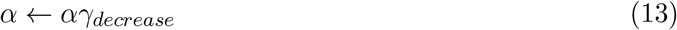

Conversely, when Δ_*improvement*_ *>* 0, we restore *α* by increasing it with a factor *γ*_*increase*_ and truncating it to *α*_*max*_:

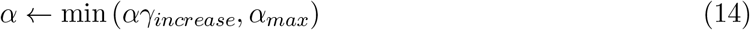

Unless otherwise specified, we set *α*_*max*_ = 0.999, *γ*_*decrease*_ = 0.6, and *γ*_*increase*_ = 1.25.

##### Note about sparse matrices and efficiency

Here, we have described the update steps (Equation 6) in terms of **D**_**r**_/**D**_**c**_ and **Q**_**r**_/**Q**_**c**_ in matrix format for the sake of brevity and clarity. While implementing the algorithm, the element-wise divide (⊘) operation can be efficiently applied on vectors and sparse matrices without ever storing the full matrices.

#### Weighted evaluation metrics

After obtaining the weighting matrix **W** using the optimization algorithm, let weighting vector 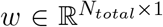 be organized in such a way that *w*_*i*_ indicates the edge weight corresponding to the *i*th prediction (after all edges are sorted based on the prediction scores of a method). Using this vector, we can compute the weighted true positives for *k* predictions as follows:

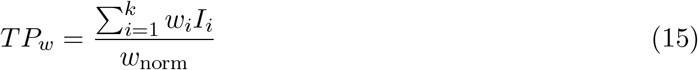

where *I*_*i*_ is an indicator variable that is equal to 1 if the ith prediction is a true positive and is equal to 0 otherwise and *w*_*norm*_ is a normalization factor:

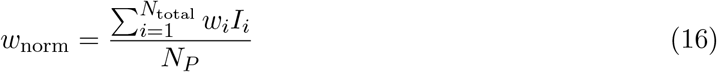

where *N*_*P*_ is the number of positive labels in the test set.

After obtaining the weighted true positives, the weighted versions of the AUPR and AUlogPR metrics are computed as described in the previous sections (this time using *TP*_*w*_ instead of *TP*).

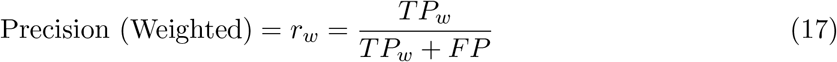

For information on how to compute credible intervals on the weighted metrics, please refer to the supplementary materials.

## Data Availability

We obtain the human protein-protein interaction network from BioGRID [35] (https://downloads.thebiogrid.org/BioGRID) and STRING [40] (http://string-db.org/cgi/download.pl). We obtain the kinase-substrate annotations from PhosphositePlus [41] (http://www.phosphosite.org/staticDownloads). We obtain the transcription factor regulatory relationships from TRRUST [42] (https://www.grnpedia.org/trrust). We obtain the drug-drug interaction dataset from Drugbank [43] and the disease-drug association datasets NDFRT and CTD [44] from BioNEV repository [1] (https://github.com/sunlab-osu/BioNEV). All materials (code and data) to reproduce the analyses and figures in the paper is available in figshare (doi: 10.6084/m9.figshare.21330429).

## Acknowledgments

This work was supported in part by the National Library Of Medicine of the National Institutes of Health under award number R01-LM012980. The content is solely the responsibility of the authors and does not necessarily represent the official views of the National Institutes of Health.

## Author Contributions

SY, KY, and MK conceived the study together. MK supervised the study, SY and KY designed and implemented the computational tools and performed data analysis together. SY was primarily responsible for data analysis and visualizations while KY was preliminary responsible for preprocessing data. All authors regularly came together and discussed the results, and collectively made decisions to drive research directions. SY and KY drafted the manuscript, MK reviewed and edited the manuscript. All authors reviewed and approved the final version of the manuscript.

## Competing Interests

The authors declare that they have no competing interests.

## Footnotes

† https://github.com/serhan-yilmaz/colipe

‡ https://yilmazs.shinyapps.io/colipe/

## Notes

https://doi.org/10.6084/m9.figshare.21330429

### Competing Interest Statement

The authors have declared no competing interest.

### Summary of Updates

Three main updates: (1) Edits for Clarity: The language was made more concise to enhance readability in the manuscript. (2) Extended Discussion and AWARE Strategies: The discussion section was expanded with additional literature, and the AWARE acronym was introduced, providing an easy to remember way of summarizing our suggestions to the community. (3) Additional Computational Experiments: New experiments were conducted on Biogrid and STRING networks in various settings. Additionally, a degree-balanced sampling strategy was introduced to reduce bias in the training of machine learning methods.

