## Supplementary Materials for "Are under-studied proteins under-represented? How to fairly evaluate link prediction algorithms in network biology"

21 S. Figure 21 - Results of STRING PPI Across-Evidence (Experimental vs. Combined) 23
22 S. Figure 22 - Comparing degree distributions STRING Experimental vs. Combined 24

#### Supplementary Methods

##### Link Prediction Algorithms - Mathematical Formulations

###### Methods based on simple scoring metrics

*Preferential Attachment:*

$$\sigma_{PA}(u,v) = \sqrt{|\Gamma(u)||\Gamma(v)|} \tag{18}$$

where $\Gamma(u)$ denotes the set containing the neighbors of $u$. In matrix form, the preferential attachment score is equal to:

$$\sigma_{PA} = \sqrt{D_r \odot D_c} \tag{19}$$

where $\odot$ indicates the element-wise (Hadamard) product and $D_r$ and $D_c$ are respectively row and column degrees in matrix form:

$$\begin{aligned} D_r(u,v) &= |\Gamma(u)| \\ D_c(u,v) &= |\Gamma(v)| \end{aligned} \tag{20}$$

*Common Neighbors:*

$$\sigma_{AA}(u,v) = |\Gamma(u) \cap \Gamma(v)| \tag{21}$$

In matrix form, this is equal to:

$$\sigma_{AA} = A^2 \tag{22}$$

where $A$ is the adjacency matrix of the network.
*Jaccard Index:*

$$\sigma_{JI}(u,v) = \frac{|\Gamma(u) \cap \Gamma(v)|}{|\Gamma(u) \cup \Gamma(v)|} \tag{23}$$

In matrix form,

$$\begin{aligned} \sigma_{JI} &= A^2 \oslash N \\ N &= D_r + D_c - A^2 \end{aligned} \tag{24}$$

where $\oslash$ indicates the element-wise (Hadamard) divide operation.

###### Higher-order paths and Network propagation based methods

*L3:* In matrix form,

$$\begin{aligned} \sigma_{L3} &= A' \times A' \times A \\ A' &= A \oslash \sqrt{D_r} \end{aligned} \tag{25}$$

*L3-Normalized (L3n):*

$$\begin{aligned} \sigma_{L3n} &= A_n^3 \\ A_n &= A \oslash D_r \end{aligned} \tag{26}$$

*von Neumann:*

$$
\begin{aligned}
A_s &= A \oslash \sqrt{D_r \odot D_c} \\
\sigma_{VN} &= \alpha A_s + \alpha^2 A_s^2 + \alpha^3 A_s^3 \dots \\
&= \sum_{i=1}^{l} \alpha^i A_s^i
\end{aligned} \tag{27}
$$

Here, we use $\alpha = 0.5$ and go up to path lengths of $l = 4$ for computational efficiency reasons.

*Random walks with restarts (RWR):*

$$
\begin{aligned}
A_n &= A \oslash D_r \\
\sigma_{RWR} &= \alpha A_n + \alpha^2 A_n^2 + \alpha^3 A_n^3 \dots \\
&= \sum_{i=1}^{l} \alpha^i A_n^i
\end{aligned} \tag{28}
$$

Similar to von-Neumann method, we use $\alpha = 0.5$ and go up to $l = 4$.

**Embedding-based methods**

In addition to these methods that compute a single score for each candidate pair, recent link prediction algorithms commonly use node embeddings to facilitate supervised learning. Node embeddings map the nodes in a network to a lower-dimensional embedding space, such that adjacent nodes are mapped to points that are close to each other in this embedding space [14]. Subsequently, using these embeddings as feature vectors and existing edges as training data, machine learning models are trained to predict new edges. We consider two embedding methods that are representative of common approaches to the computation of node embeddings.

**Deepwalk (Random walk based node embedding):** Deepwalk [13] uses random walks to generate a list of paths in the network as its corpus and then uses Word2Vec [47], a natural language processing algorithm for word embedding, to compute node embeddings by treating the list of paths as text and nodes as words. In our experiments, we use the implementation used in BioNEV [1] repository (which is based on OpenNE [48]) for all embedding methods.

**LINE (Neural network based embedding):** LINE [36] is one of the earliest algorithms to incorporate neural networks into the computation of node embeddings. It uses a single layer MLP to estimate first and second order proximity of nodes and produces the embedding vectors using a variational auto-encoder.

Unless otherwise specified, we use the default value of 128 in the OpenNE implementation as the embedding dimension (i.e., number of embeddings) for both embedding methods.

In addition to the above two, we consider a version of deepwalk (***Deepwalk-withdegree***) where the node degree information is appended as an additional dimension to the embedding matrix.

**Logistic regression as prediction model:** For each of three embedding approaches described above, we train a logistic regression using the embeddings as features. For each embedding dimension $x^{(i)}$, we add the following three terms to the logistic regression model corresponding to the prediction for edge (u,v):

$$
\text{logit}(Y_{uv}) \propto \sum_i \left( \beta_r^{(i)} x_u^{(i)} + \beta_c^{(i)} x_v^{(i)} + \beta_{rc}^{(i)} x_u^{(i)} x_v^{(i)} \right) \tag{29}
$$

While training the model, to ensure a balanced training set, we randomly sample the edges with negative labels (i.e., not in the training set) to have the same size as the edges with positive labels.

#### Evaluation Metrics for Prediction Performance

*Precision and Scaled Precision:*

$$\text{Precision} = r = \frac{TP}{TP + FP} \tag{30}$$

where $TP$ denotes the number of true positives, and $FP$ the number of false positives. The expected precision for random predictions is equal to the prevalence of positive labels:

$$E[\text{Precision}] = \text{Prevalence} = \frac{N_P}{N_{\text{total}}} \tag{31}$$

where $N_P$ is the number of positive labels and $N_{\text{total}}$ is the total number of edges that are to be predicted (approximately $O(N^2)$).

$$\text{Scaled Precision} = \frac{\text{Precision}}{E[\text{Precision}]} = \frac{TP}{TP + FP}\frac{N_{\text{total}}}{N_P} \tag{32}$$

*Recall:*

$$\text{Recall} = \frac{TP}{N_P} \tag{33}$$

##### Computing the area under precision-recall curve (AUPR)

To compute the area under the precision-recall (PR) curve, we use numerical integration. Suppose we have $m$ measurement points. Let $TP_i$ denote the number of true positives, $FP_i$ the number of false positives, $N_i = TP_i + FP_i$ the number of predictions, $X_i$ the recall, and $Y_i$ the precision corresponding to the $i$th measurement point. In general, $m$ is less than $N_{total}$ since edges having the same prediction score (e.g., because the link prediction method uses discrete scoring like common neighbors) correspond to a single measurement point. Also, without loss of generality, consider that the first point is the $TP = 0$ and $FP = 0$ point and all points are sorted by the number of predictions ($TP_i + FP_i$) in ascending order. With these in mind, we compute the area under precision-recall curve through numerical integration as follows:

$$\text{AUPR} = \frac{\sum_{i=1}^{m-1} \Delta X_i f Y_i}{\sum_{i=1}^{m-1} \Delta X_i} \tag{34}$$

where $\Delta X_i$ is the gap between two consecutive points:

$$\Delta X_i = |X_{i+1} - X_i| \tag{35}$$

Whereas, $fY_i$ is an interpolating function that returns the normalized area under two consecutive points $Y_i$ and $Y_{i+1}$ (thus, it is a type of averaging for two given points and is always between $[Y_i, Y_{i+1}]$). For example, a simple function for this purpose can be $\frac{Y_i + Y_{i+1}}{2}$ (interpolating the precision values linearly). However, this type of interpolation suffers from inaccuracy when there are large gaps between two consecutive points $X_i$ and $X_{i+1}$, which is particularly relevant for link prediction methods with discrete scoring. To demonstrate the inaccuracy, suppose the first point is at 1/1 ($TP = 1$, $FP = 0$) with precision 1 and the next point is in 100/10000 ($TP = 100$, $FP = 9900$) with precision 0.01. In this example, although linear interpolation suggests that the average precision would be ≈0.5, observing one TP in the beginning hardly gives any evidence that the precision will be ≈0.5 at the $TP = 5000$ point. To overcome this type of inaccuracy, we use an interpolation function tailored for the precision-recall curve detailed below:

**Interpolating the curve during numerical integration for computing the area under**

For the intermediate points between two consecutive points $X_i$ and $X_{i+1}$, we assume that both the true positives and false positives are scaled linearly:

$$\begin{aligned} TP_x &= TP_i + x(TP_{i+1} - TP_i) \\ FP_x &= FP_i + x(FP_{i+1} - FP_i) \end{aligned} \tag{36}$$

where $x$ is a normalized variable between [0, 1] indicating which endpoint the point is closest to (e.g., 1 indicates the point is right on the i+1th point). Thus, the precision for the intermediate points is given by the ratio $r_i(x)$:

$$\begin{aligned} r_i(x) &= \frac{TP_i + x(TP_{i+1} - TP_i)}{TP_i + x(TP_{i+1} - TP_i) + FP_i + x(FP_{i+1} - FP_i)} \\ &= \frac{TP_i + x(TP_{i+1} - TP_i)}{N_i + x(N_{i+1} - N_i)} \end{aligned} \tag{37}$$

To compute the area under this curve (denoted $fY$), we need the integral:

$$fY_i = \int_0^1 r_i(x)dx = \int_0^1 \frac{TP_i + x(TP_{i+1} - TP_i)}{N_i + x(N_{i+1} - N_i)}dx \tag{38}$$

Solving this integral gives:

$$fY_i = \frac{(TP_i N_{i+1} - TP_{i+1} N_i) \log\left(\frac{N_{i+1}}{N_i}\right) + (N_{i+1} - N_i)(TP_{i+1} - TP_i)}{(N_{i+1} - N_i)^2} \tag{39}$$

Thus, we use the above function for interpolating while computing the area under the PR curve. To give some insight into what this function results in: For the example before (one point at $TP/N$ = 1/1 while the other is at 100/10000), this integral results in $\approx$ 0.011 precision which is much closer to the latter point (as it should be).

Note that, although this integral (and Equation 39) is not defined at $N_i = 0$ point (since precision is not defined at 0 predictions), the limit from above converges to $\frac{TP_{i+1}}{N_{i+1}} = Y_{i+1}$. Thus, as the first interpolated area, we use:

$$fY_1 = Y_2 \tag{40}$$

where $Y_2$ (i.e., the second point) corresponds to the first measured precision value (since the 0/0 point is specified in the $i = 1$th point in this notation).

Overall, this interpolation is helpful for reducing the inaccuracy when there are large gaps in between, which is particularly relevant for methods with discrete scoring or for computing the area under the PR curve in logarithmic scale.

**Computing credible intervals for the variance in estimation**

We follow a Bayesian approach to estimate the expected variance in the evaluation metrics (e.g., precision and AUPR). Our view here is akin to the ”checking whether a coin is fair” problem. We assume that there is an unknown, but fixed probability $r$ (corresponding to precision). Based on this probability, we suppose that we have made $k$ trials (corresponding to predictions) and observed $TP$ number of hits and $FP$ number of misses. Now, we ask the question ”Based on these observations, what can we say about the posterior probability of the ratio $r$?”

If we assume uniform prior (i.e., all $r$ values in [0, 1] are equally likely), the answer to the above question is specified by the beta distribution:

$$r \sim \text{Beta}(TP + 1, FP + 1) \tag{41}$$

Thus, we obtain the distribution for the posterior probability of the ratio $r$ (i.e., precision) after $k$ predictions. Based on this distribution, we can easily construct a credible interval containing the 95% of the variance using the inverse cumulative distribution function $\text{Beta}^{-1}$. Note that, in general, there is not a single credible interval unique to a given posterior distribution. Thus, among the alternatives, we choose the equal-tailed interval where the probability of being below the interval is as likely as being above it.

This process gives us a 95% interval for the precision at fixed number of predictions $k$ point. To obtain 95% intervals for the area under metrics (AUPR and AUlogPR), we simply construct the intervals for all $k$ points and compute the area under the precision-recall curves formed by the maximum/minimum bounds.

##### Computing credible intervals for the weighted metrics

In the previous section *"Computing credible intervals for the variance in estimation"*, we obtained the posterior distribution of the ratio $r$ corresponding to unweighted precision (Equation 41). Here, we will transform this for the weighted precision. For this purpose, we start by defining a weighting factor $w_f$ equal to the ratio of weighted and unweighted true positives:

$$w_f = \frac{TP_w}{TP} \tag{42}$$

Using this, we can write the equation for weighted precision in terms of the unweighted ratio $r$:

$$\begin{aligned} r_w = \frac{TP_w}{TP_w + FP} &= \frac{w_f TP}{w_f TP + FP} \\ = \frac{\dfrac{w_f TP}{TP + FP}}{\dfrac{w_f TP + FP}{TP + FP}} &= \frac{w_f r}{(w_f - 1)r + 1} \end{aligned} \tag{43}$$

Thus, we transform the distribution given in Equation 41 according to the above equation to obtain the posterior distribution of the weighted ratio $r_w$. After that, we compute the 95% credible intervals as detailed before.

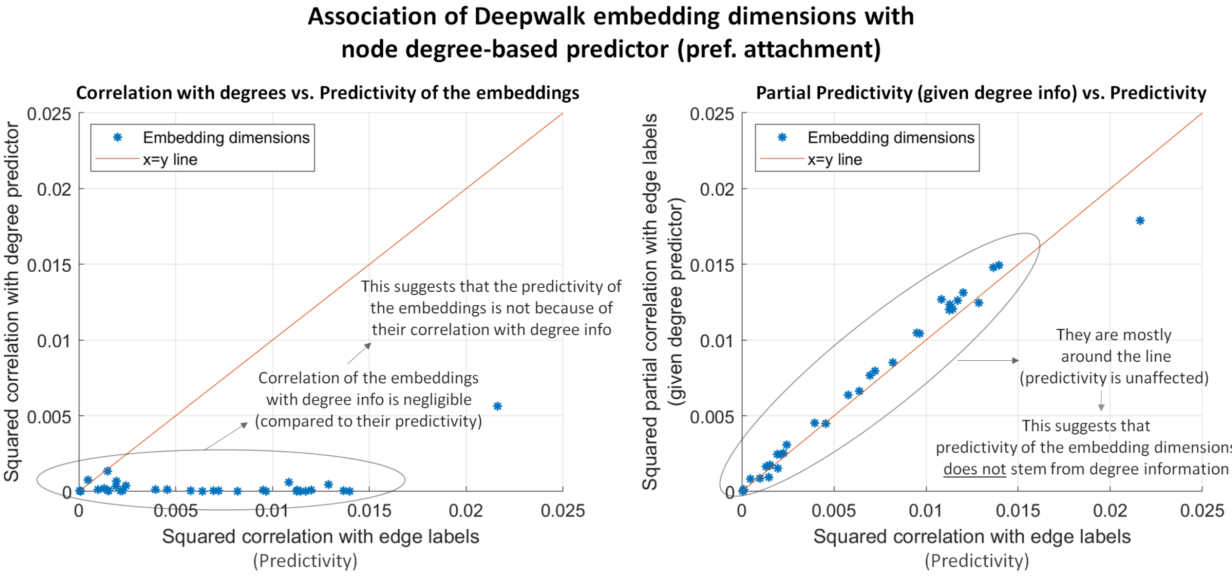

**S. Figure 1: Investigating the embedding dimensions of Deepwalk in terms of their association with node degrees.** The analysis suggests that the embeddings of Deepwalk does not depend on the degree information.

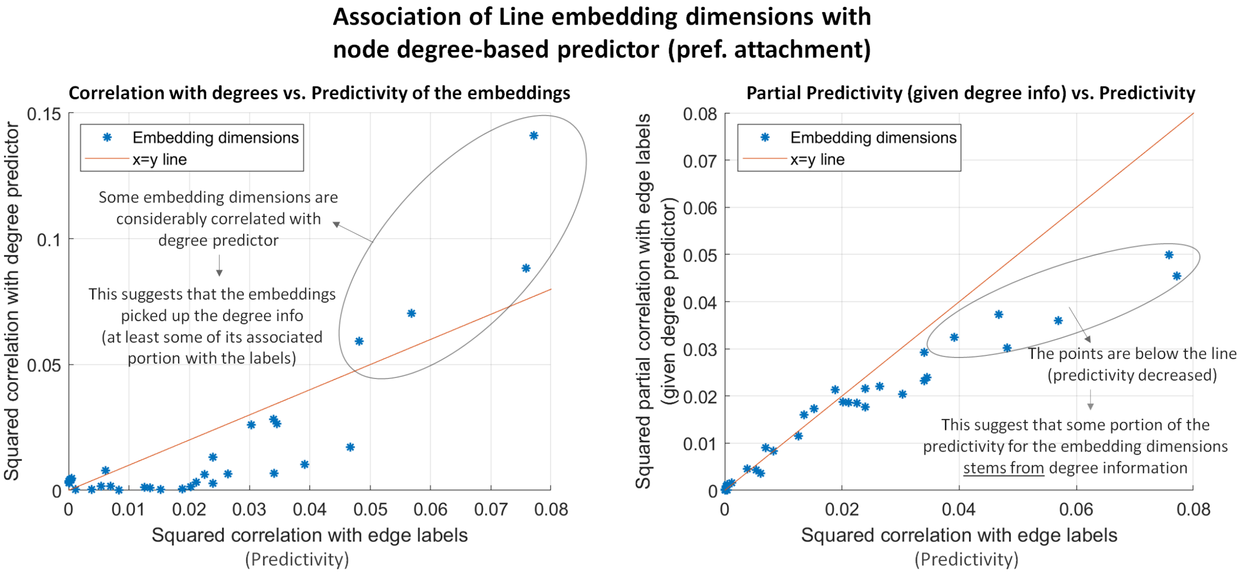

**S. Figure 2: Investigating the embedding dimensions of Line in terms of their association with node degrees.** The analysis suggests that Line embeddings picked up the node degree information and the predictivity of some of its embeddings dimensions stems from their correlation with node degrees.

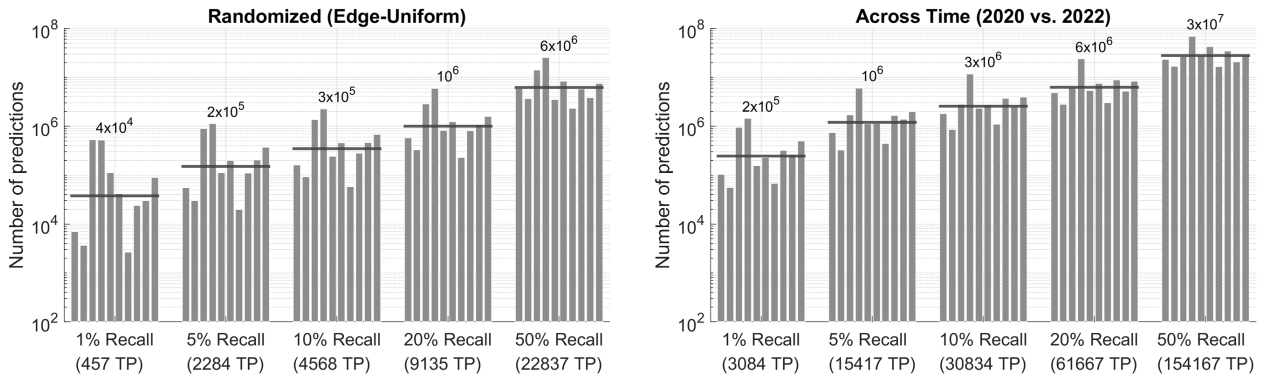

**S. Figure 3: Number of predictions required to reach a particular recall threshold for Biogrid PPI predictions.** (Left) The randomized (edge-uniform) evaluation. (Right) The across time evaluation setting. For both panels, the bars represent different link prediction algorithms. The horizontal lines and the numbers on the top indicate the geometric average of the number of predictions for each recall threshold.

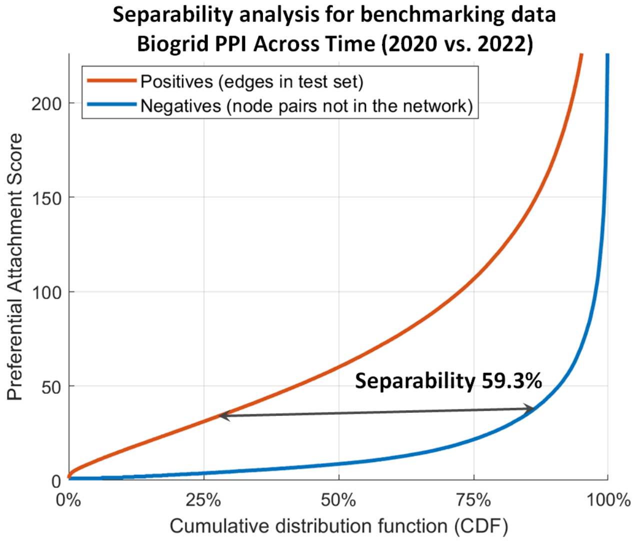

**S. Figure 4: Separability analysis investigating the informedness of node degree information for Biogrid PPI across time (2020 vs. 2022) setting.**

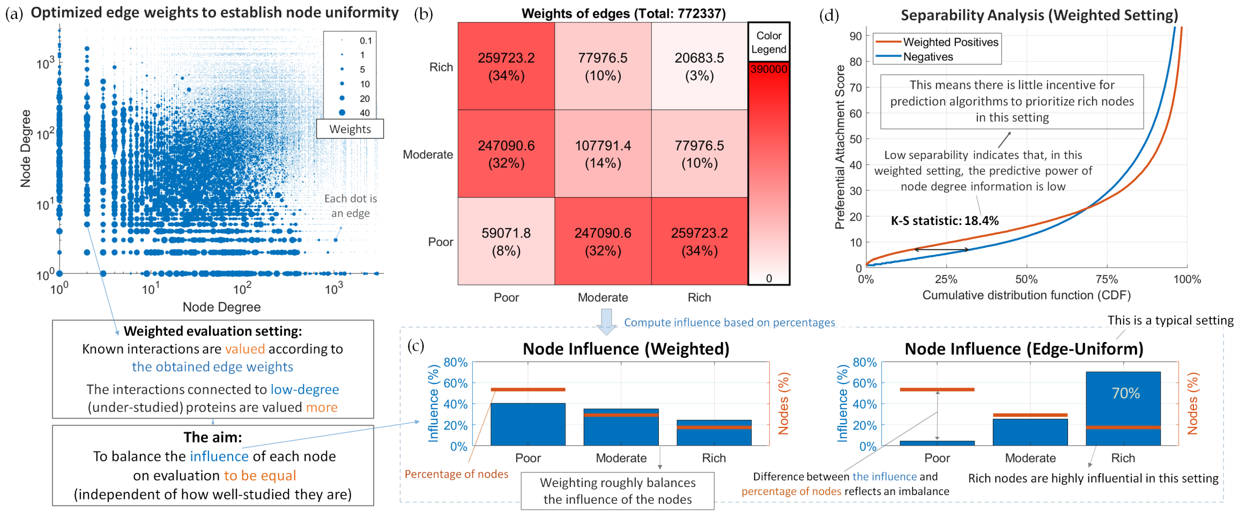

**S. Figure 5: Mitigating degree bias in the evaluation of link prediction algorithms by assigning weights to edges during evaluation. Assignment of optimized edge weights establishes node-uniformity and balances the influence of nodes on evaluation.** (a) Visualization of the optimized edge weights with respect to the degrees of incident nodes. The size of each point reflects the assigned weight of the corresponding edge. (b) The total weight of the edges by node category. (c) Influence of the nodes on evaluation (shown as bars) with respect to the node categories for weighted (node-uniform) and unweighted (edge-uniform) settings. (c) Separability analysis for the weighted (node-uniform) evaluation setting.

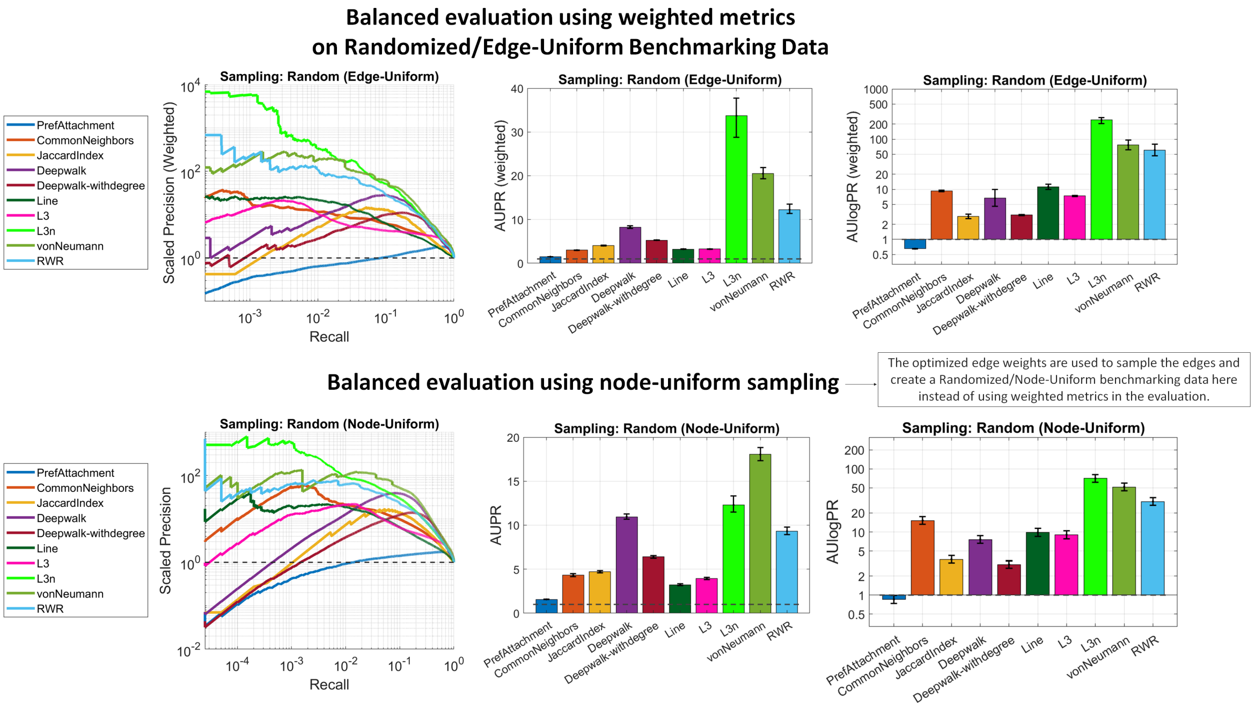

**S. Figure 6: Balanced/Weighted evaluation results on randomized (sampled) benchmarking data for Biogrid PPI predictions.** (Top) Balanced evaluation using weighted metrics. (Bottom) Balanced evaluation via node-uniform sampling (using the weights as sampling probabilities)

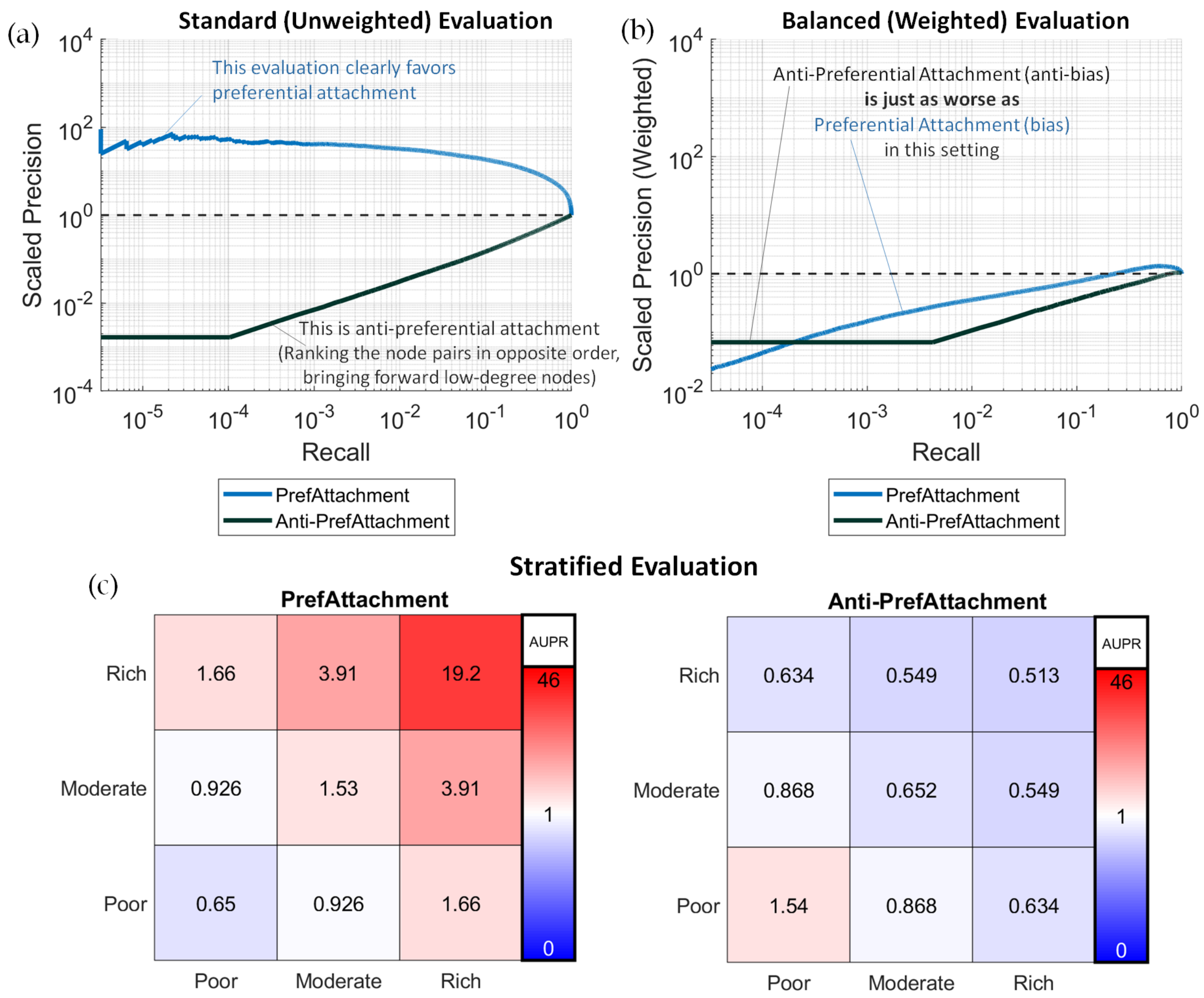

**S. Figure 7: Comparison of preferential attachment (biased baseline) and anti-preferential attachment (anti-biased baseline) in different evaluation settings on Biogrid PPI predictions**. Across-time (2020 vs. 2022) snapshots of the network are used as the benchmarking data (i.e., train/test splits) in this analysis. (a & b) Precision-Recall curves for the preferential attachment and anti-preferential attachment models respectively for standard (unweighted) and balanced (weighted) evaluation settings. (c) Stratified performance analysis results for preferential attachment and anti-preferential attachment algorithms. Each cell indicates the prediction performance of the algorithms for the corresponding edge category (e.g., for Poor-Rich category, only the edges that are between poor and rich nodes are included in the test set).

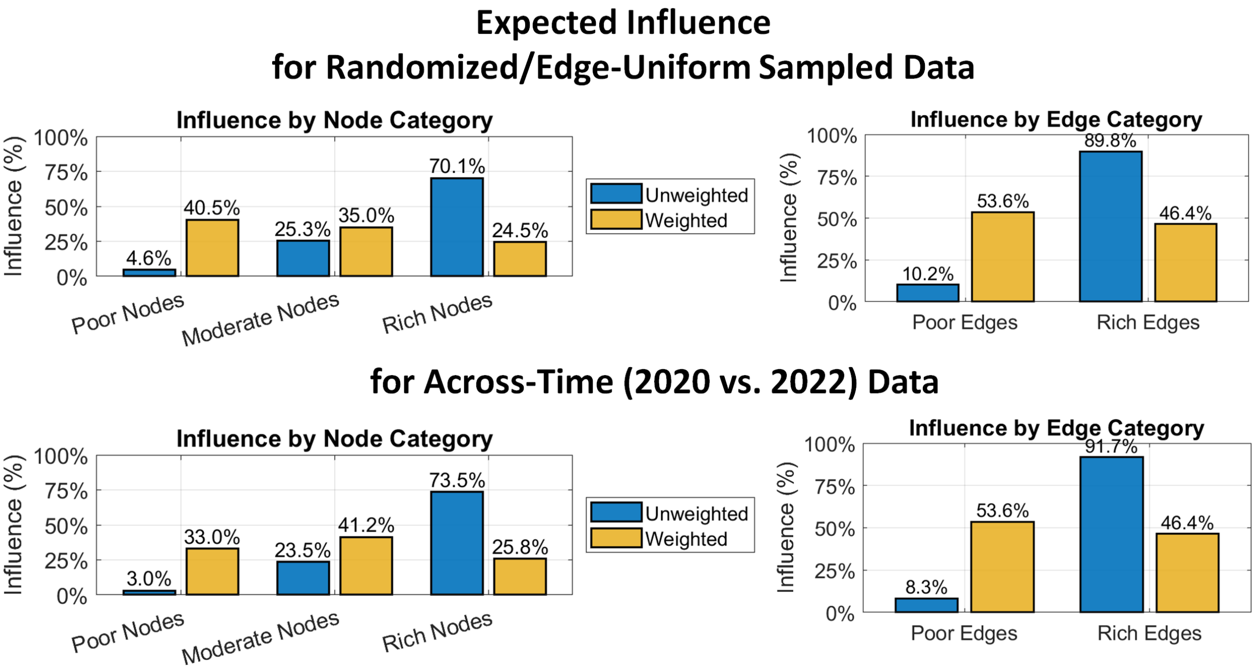

**S. Figure 8: Expected influence for different categories of nodes or edges based on node degrees for randomized/edge-uniform and across-time bencharking data.**

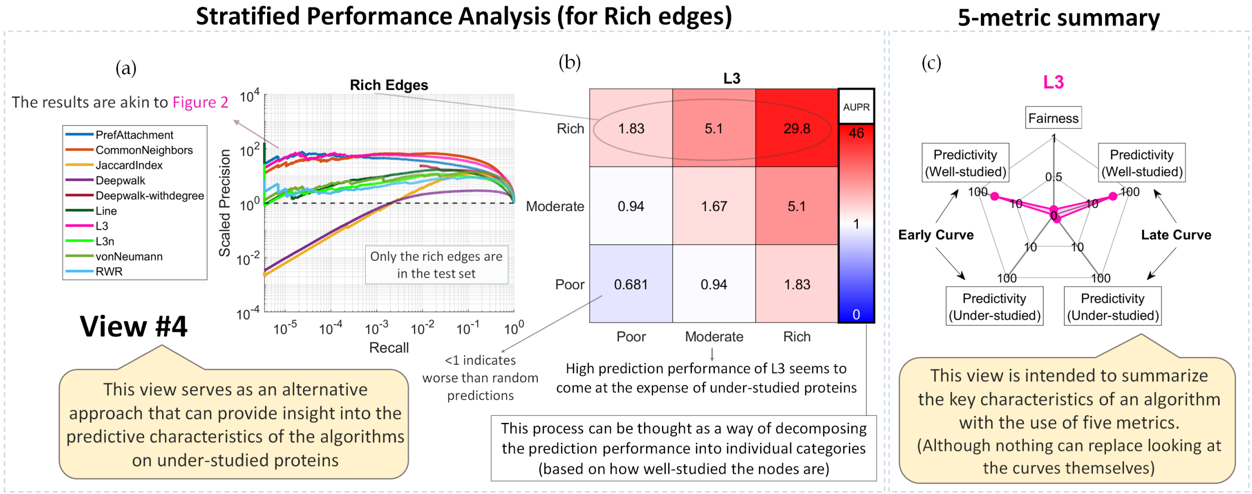

**S. Figure 9: Stratified performance analysis for Rich edges connnected to well-studied nodes and the 5-metric summary for the best performing method on rich edges.** (a) Precision-Recall performance curves for rich edges in log-log scale. (b) Late curve prediction performance (AUPR) stratified by node categories for L3 algorithm. (c) 5-metric summary for L3.

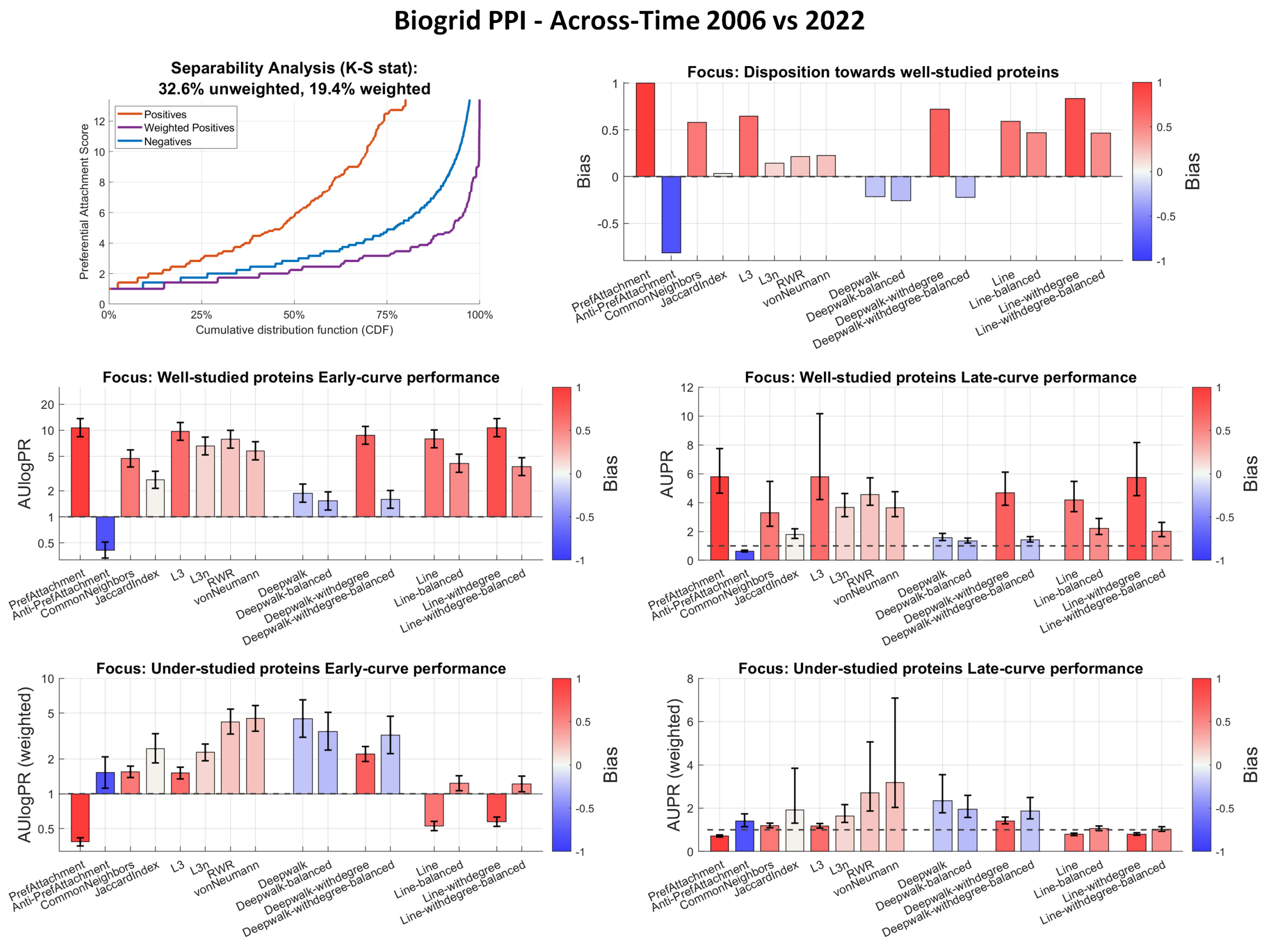

**S. Figure 10: Results of bias-aware evaluation on Biogrid PPI Across-Time (2006 vs. 2022).** (Top Left) Separability analysis. (Remaining Panels) Evaluation results for each of the five metrics are shown with bar plots: Bias score (based on similarity with preferential attachment), AUPR, AUlogPR, weighted AUPR and weighted AUlogPR to focus on under-studied proteins. In each panel, the coloring is done according to the bias scores.

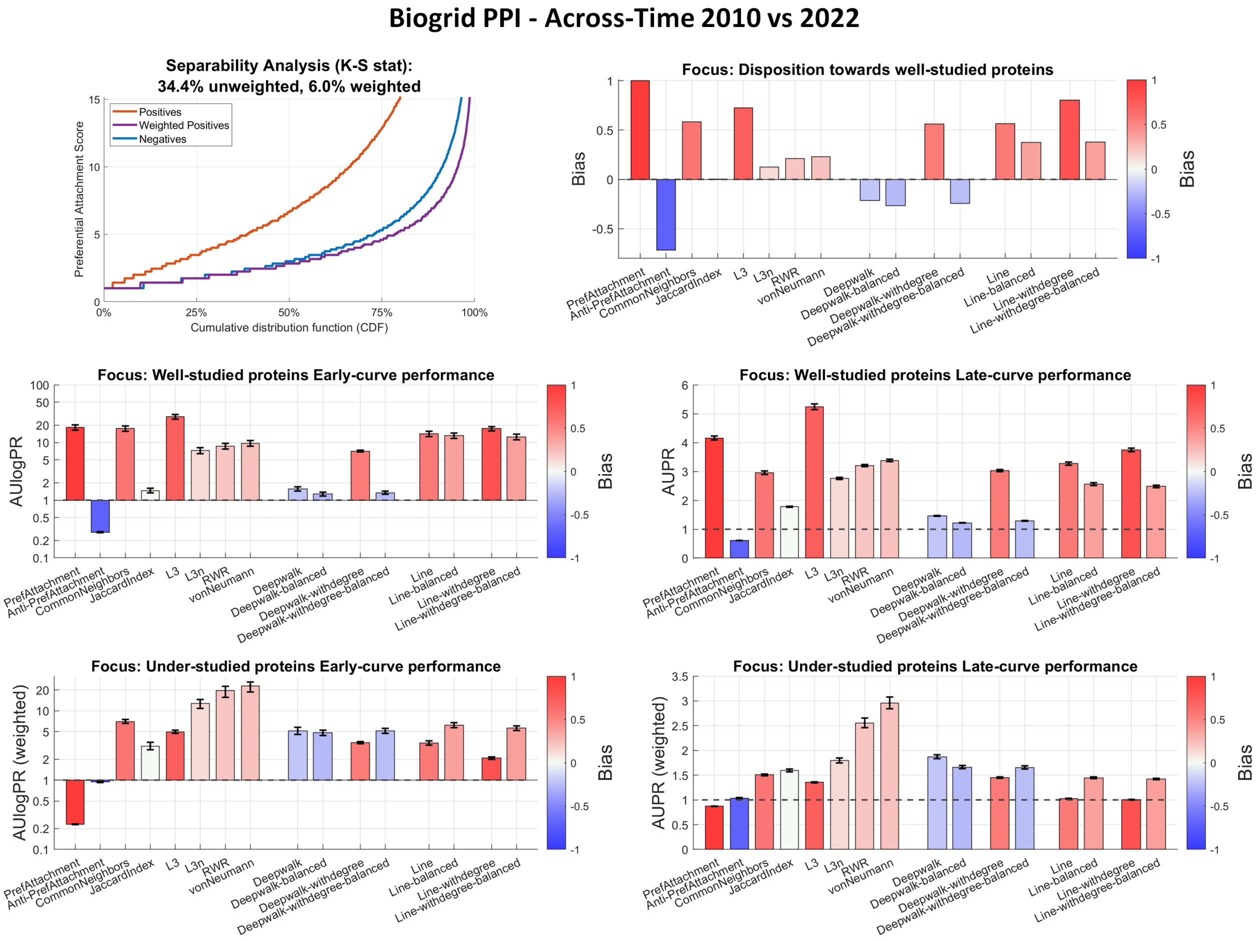

**S. Figure 11: Results of bias-aware evaluation on Biogrid PPI Across-Time (2010 vs. 2022).** (Top Left) Separability analysis. (Remaining Panels) Evaluation results for each of the five metrics are shown with bar plots: Bias score (based on similarity with preferential attachment), AUPR, AUlogPR, weighted AUPR and weighted AUlogPR to focus on under-studied proteins. In each panel, the coloring is done according to the bias scores.

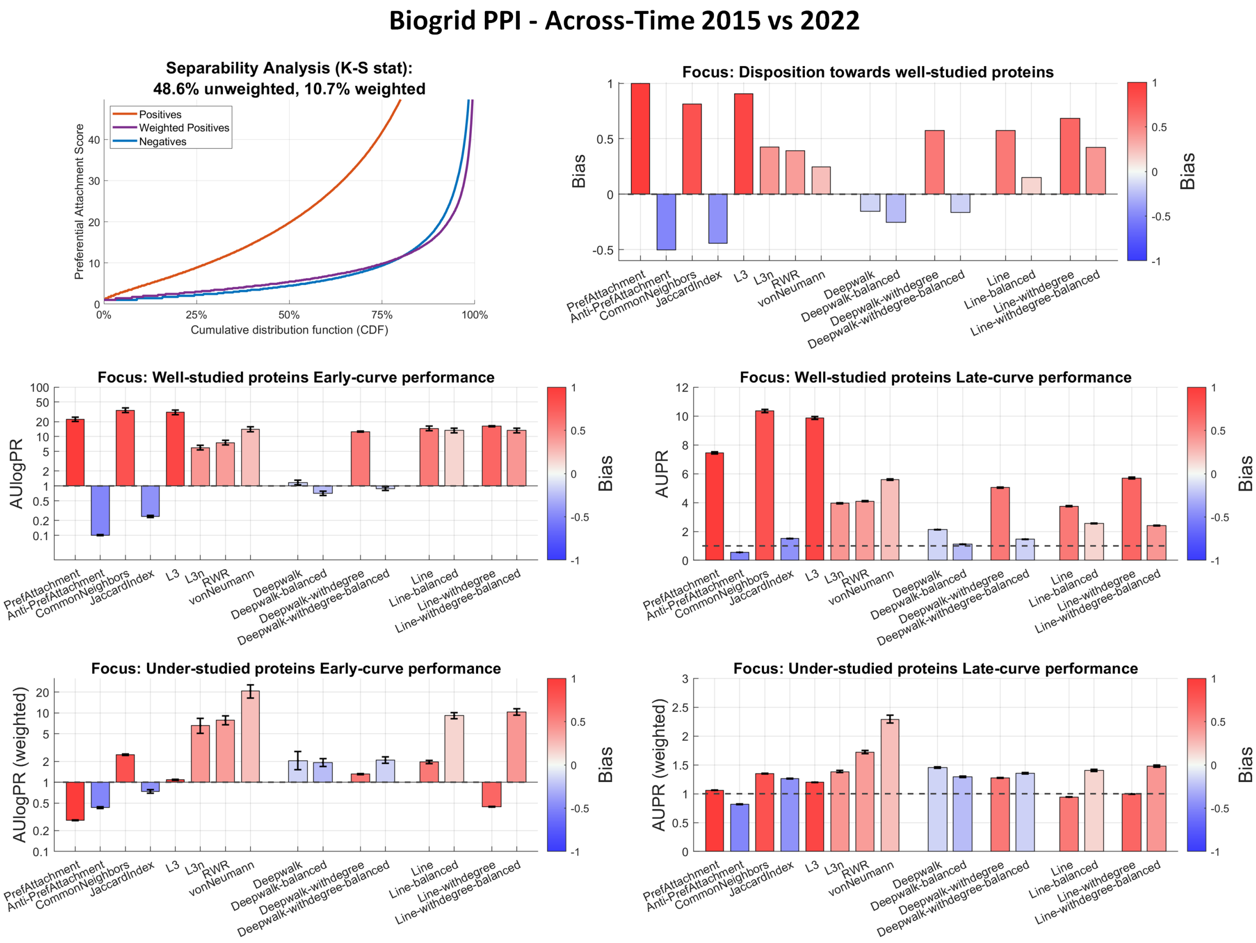

**S. Figure 12: Results of bias-aware evaluation on Biogrid PPI Across-Time (2015 vs. 2022).** (Top Left) Separability analysis. (Remaining Panels) Evaluation results for each of the five metrics are shown with bar plots: Bias score (based on similarity with preferential attachment), AUPR, AUlogPR, weighted AUPR and weighted AUlogPR to focus on under-studied proteins. In each panel, the coloring is done according to the bias scores.

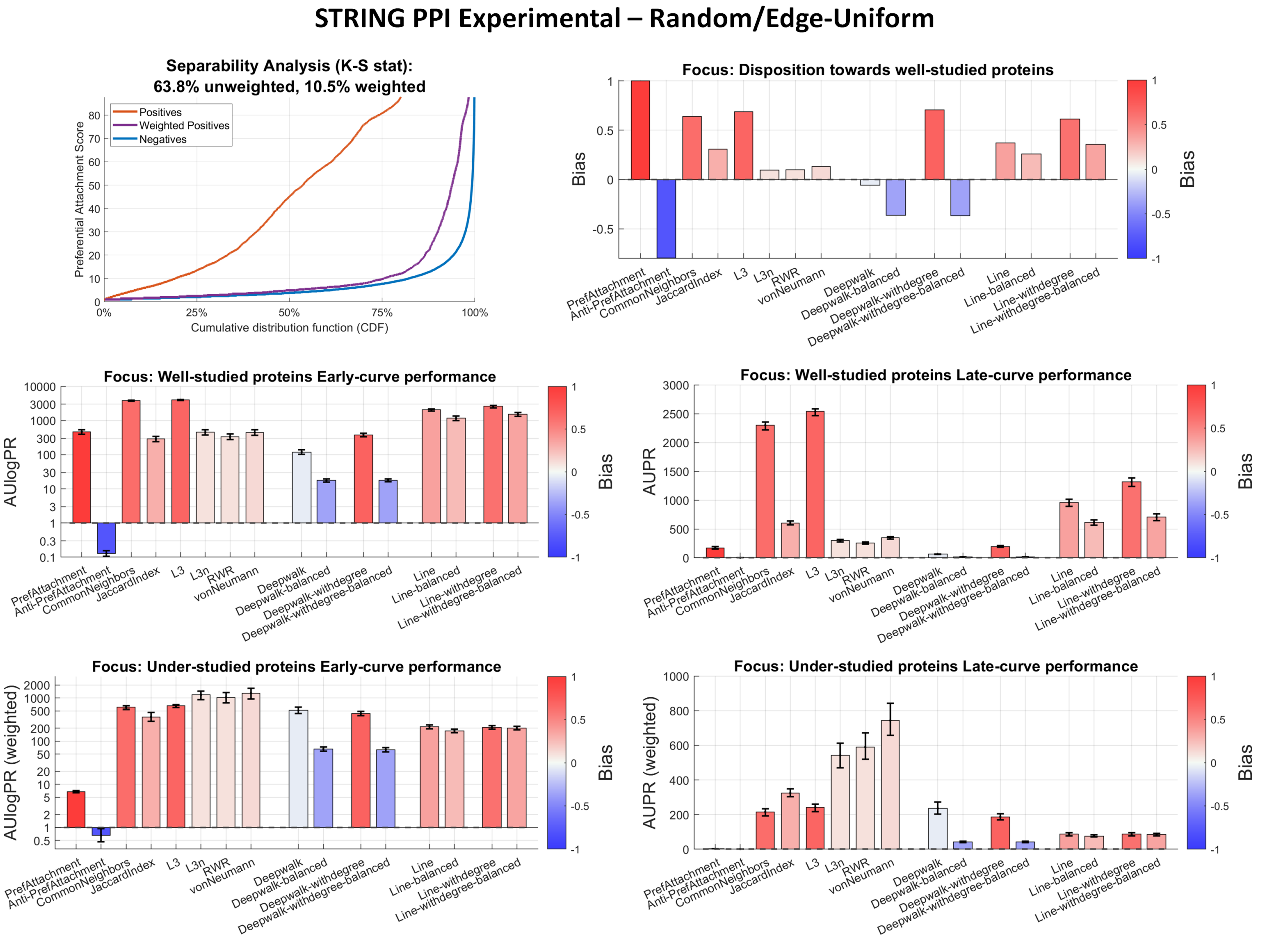

**S. Figure 13: Results of bias-aware evaluation on STRING PPI experimental sub-network Random/Edge-Uniform setting.** (Top Left) Separability analysis. (Remaining Panels) Evaluation results for each of the five metrics are shown with bar plots: Bias score (based on similarity with preferential attachment), AUPR, AUlogPR, weighted AUPR and weighted AUlogPR to focus on under-studied proteins. In each panel, the coloring is done according to the bias scores.

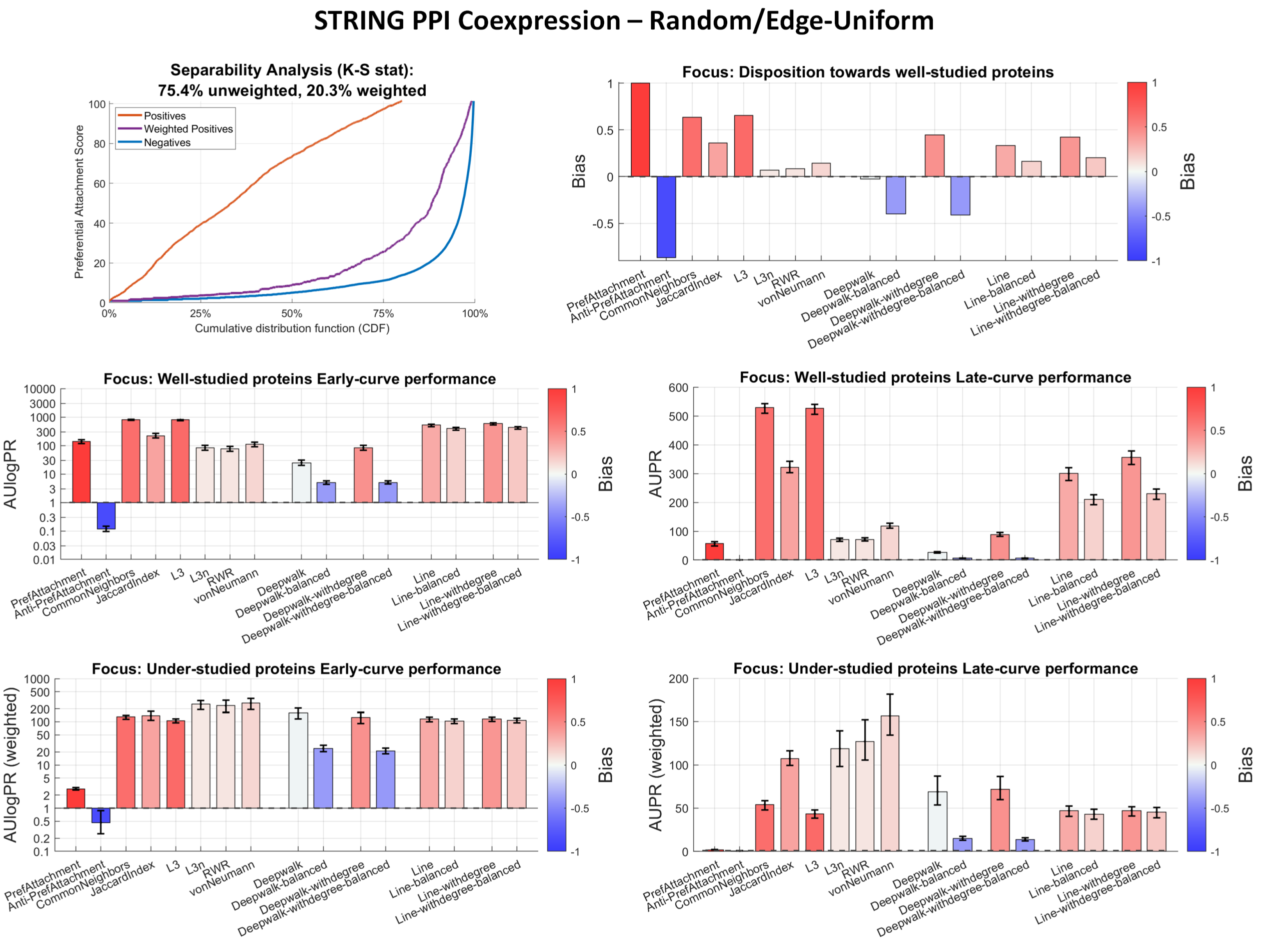

**S. Figure 14: Results of bias-aware evaluation on STRING PPI coexpression sub-network Random/Edge-Uniform setting.** (Top Left) Separability analysis. (Remaining Panels) Evaluation results for each of the five metrics are shown with bar plots: Bias score (based on similarity with preferential attachment), AUPR, AUlogPR, weighted AUPR and weighted AUlogPR to focus on under-studied proteins. In each panel, the coloring is done according to the bias scores.

#### STRING PPI Textmining – Random/Edge-Uniform

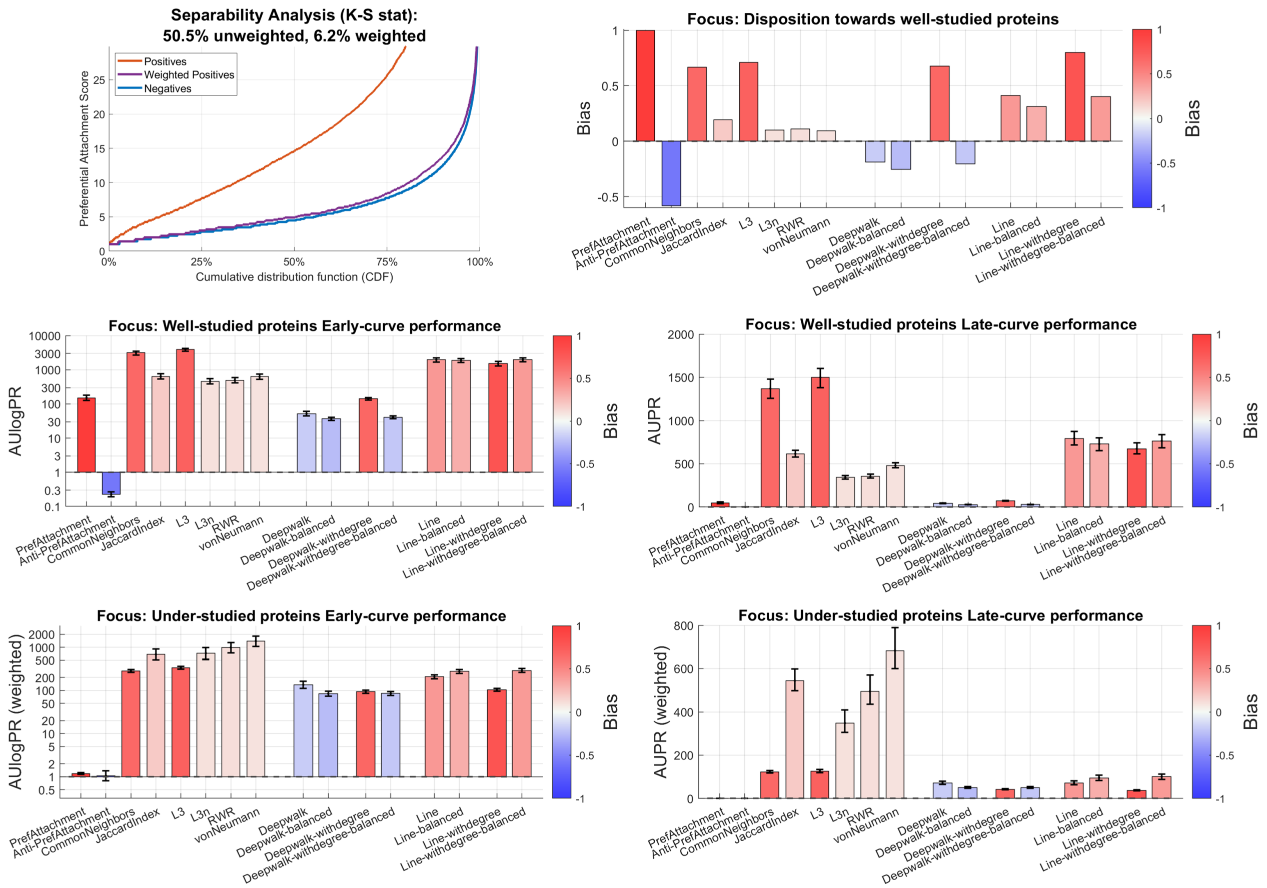

**S. Figure 15: Results of bias-aware evaluation on STRING PPI textmining sub-network Random/Edge-Uniform setting.** (Top Left) Separability analysis. (Remaining Panels) Evaluation results for each of the five metrics are shown with bar plots: Bias score (based on similarity with preferential attachment), AUPR, AUlogPR, weighted AUPR and weighted AUlogPR to focus on under-studied proteins. In each panel, the coloring is done according to the bias scores.

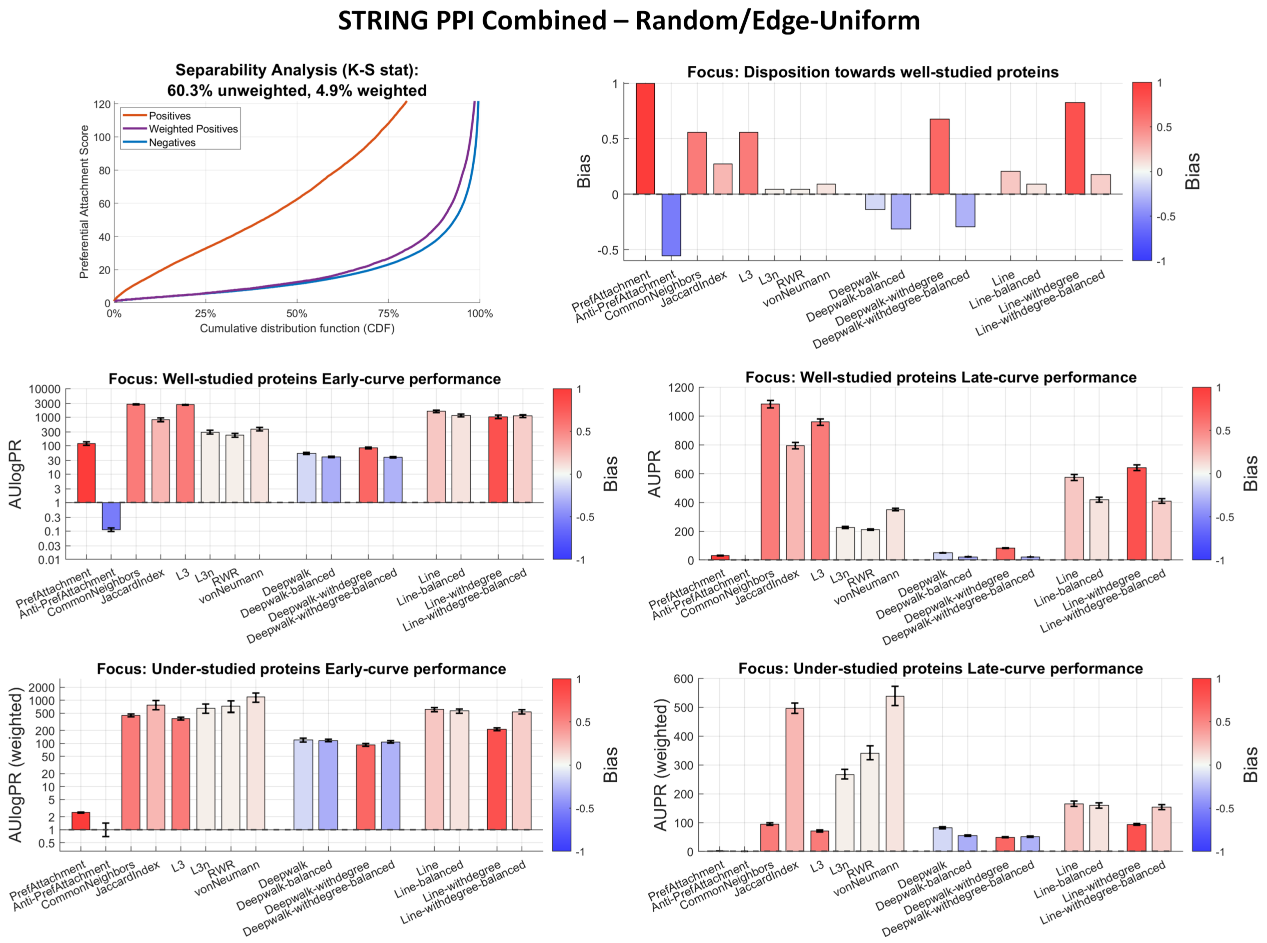

**S. Figure 16: Results of bias-aware evaluation on STRING PPI combined network Random/Edge-Uniform setting.** (Top Left) Separability analysis. (Remaining Panels) Evaluation results for each of the five metrics are shown with bar plots: Bias score (based on similarity with preferential attachment), AUPR, AUlogPR, weighted AUPR and weighted AUlogPR to focus on under-studied proteins. In each panel, the coloring is done according to the bias scores.

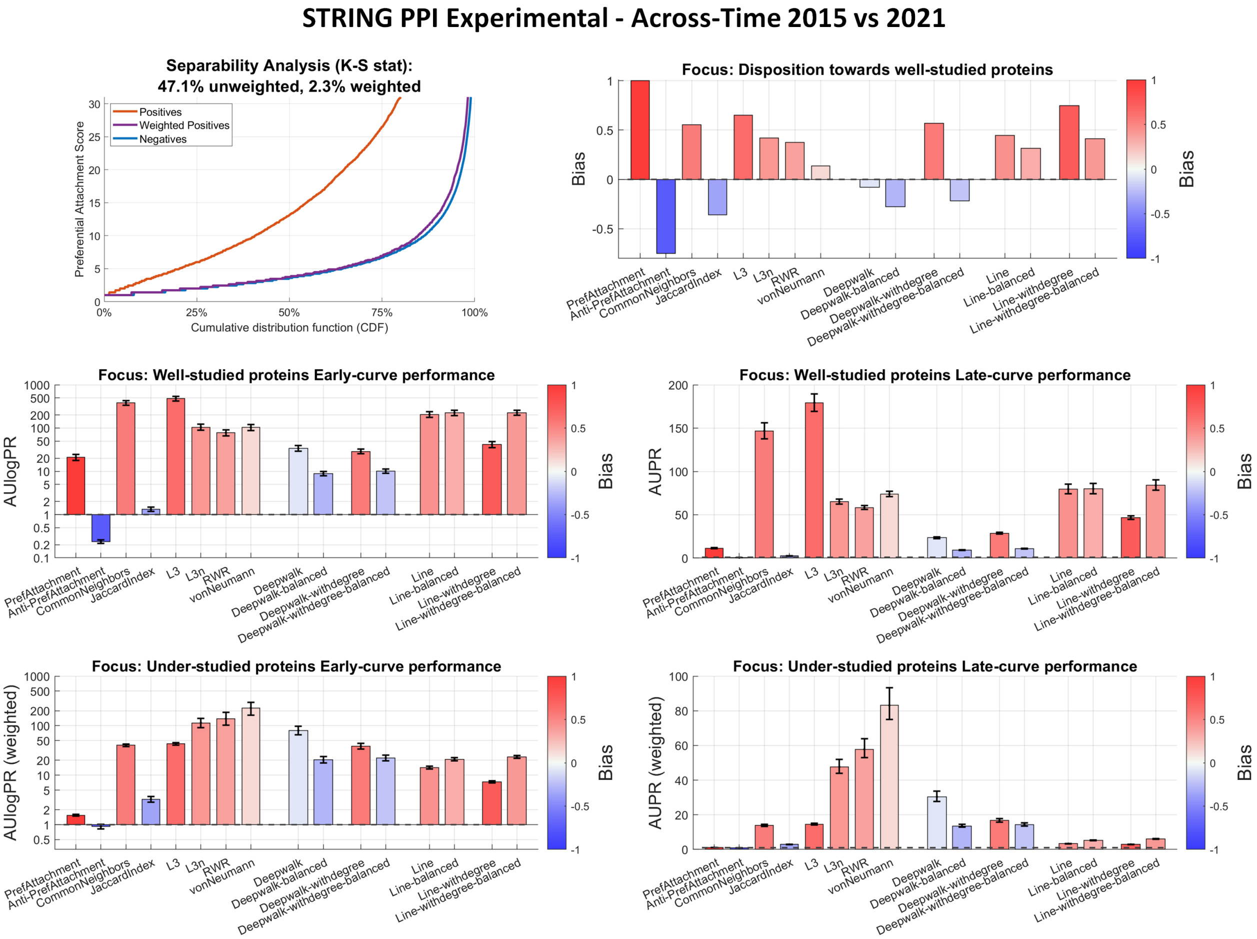

**S. Figure 17: Results of bias-aware evaluation on STRING PPI experimental sub-network Across-Time (2015 vs. 2021) setting.** (Top Left) Separability analysis. (Remaining Panels) Evaluation results for each of the five metrics are shown with bar plots: Bias score (based on similarity with preferential attachment), AUPR, AUlogPR, weighted AUPR and weighted AUlogPR to focus on under-studied proteins. In each panel, the coloring is done according to the bias scores.

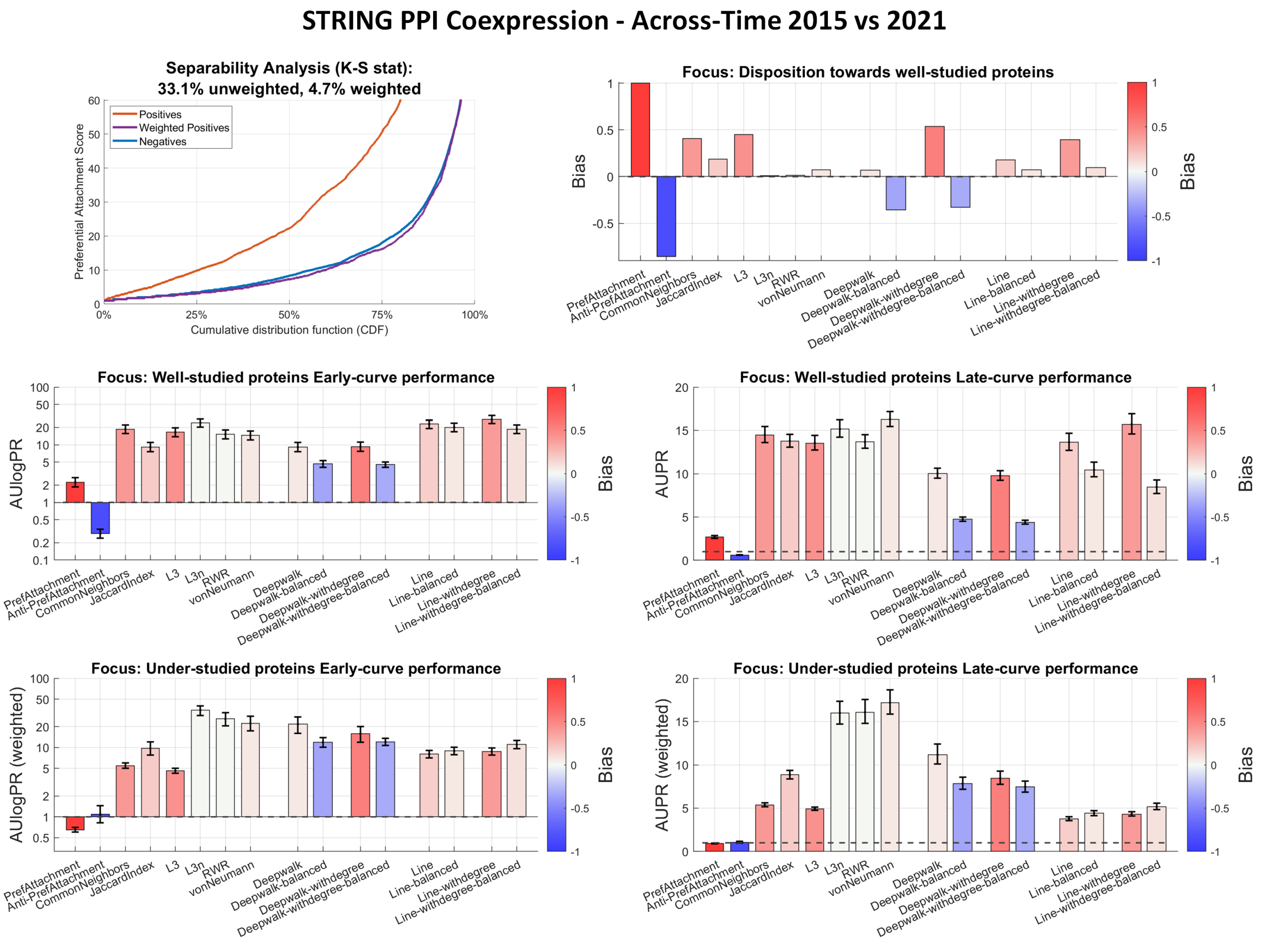

**S. Figure 18: Results of bias-aware evaluation on STRING PPI coexpression sub-network Across-Time (2015 vs. 2021) setting.** (Top Left) Separability analysis. (Remaining Panels) Evaluation results for each of the five metrics are shown with bar plots: Bias score (based on similarity with preferential attachment), AUPR, AUlogPR, weighted AUPR and weighted AUlogPR to focus on under-studied proteins. In each panel, the coloring is done according to the bias scores.

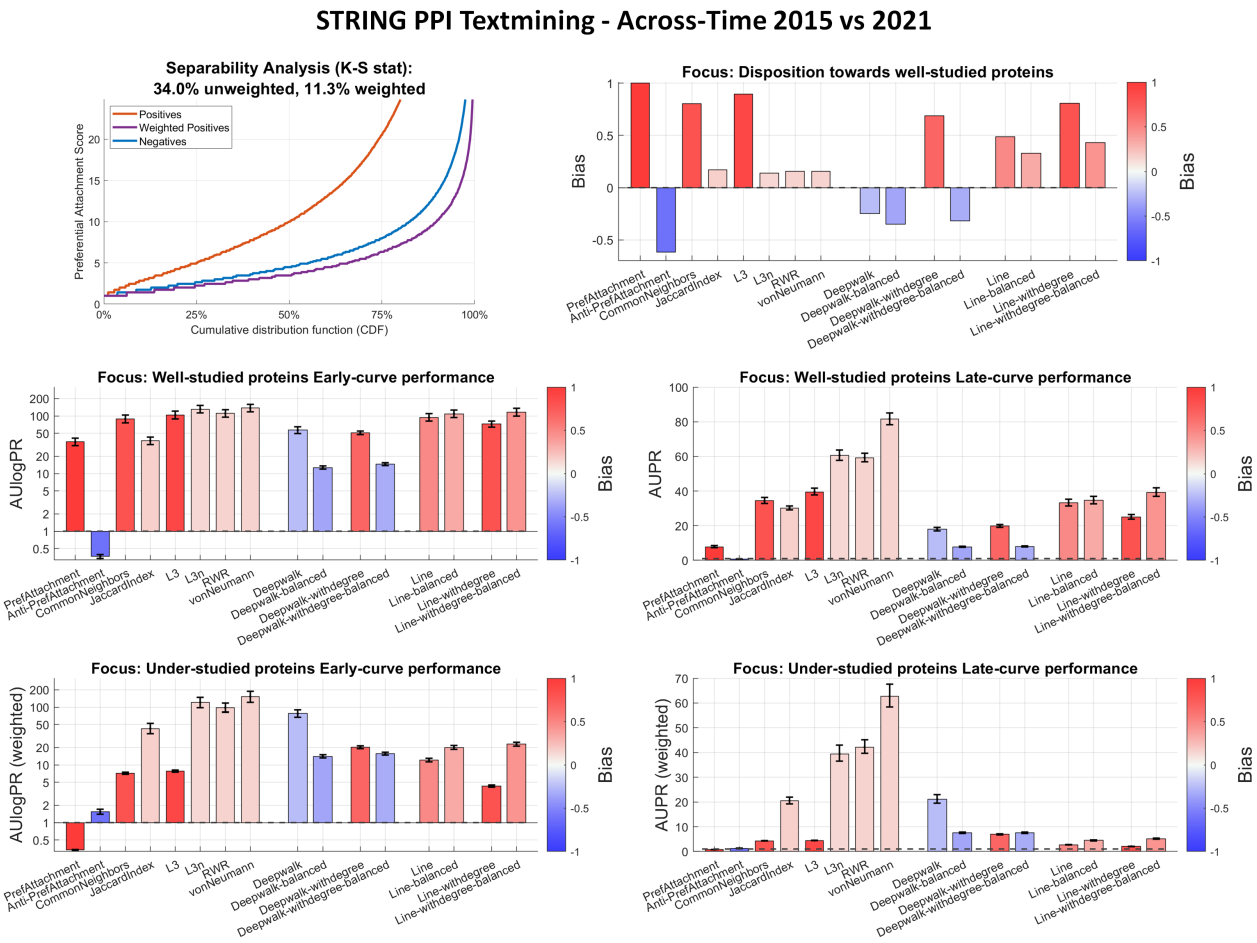

**S. Figure 19: Results of bias-aware evaluation on STRING PPI textmining sub-network Across-Time (2015 vs. 2021) setting.** (Top Left) Separability analysis. (Remaining Panels) Evaluation results for each of the five metrics are shown with bar plots: Bias score (based on similarity with preferential attachment), AUPR, AUlogPR, weighted AUPR and weighted AUlogPR to focus on under-studied proteins. In each panel, the coloring is done according to the bias scores.

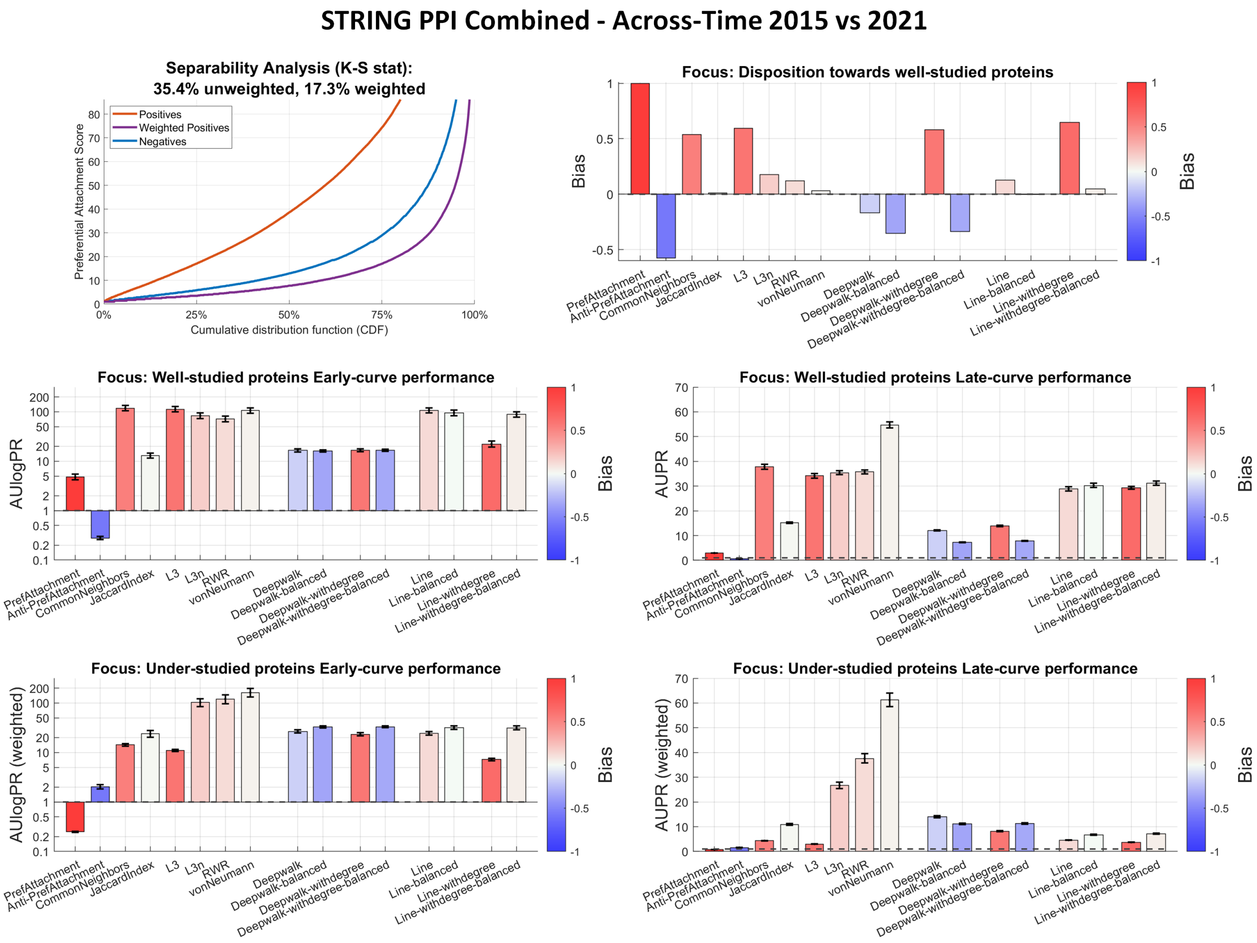

**S. Figure 20: Results of bias-aware evaluation on STRING PPI combined network Across-Time (2015 vs. 2021) setting.** (Top Left) Separability analysis. (Remaining Panels) Evaluation results for each of the five metrics are shown with bar plots: Bias score (based on similarity with preferential attachment), AUPR, AUlogPR, weighted AUPR and weighted AUlogPR to focus on under-studied proteins. In each panel, the coloring is done according to the bias scores.

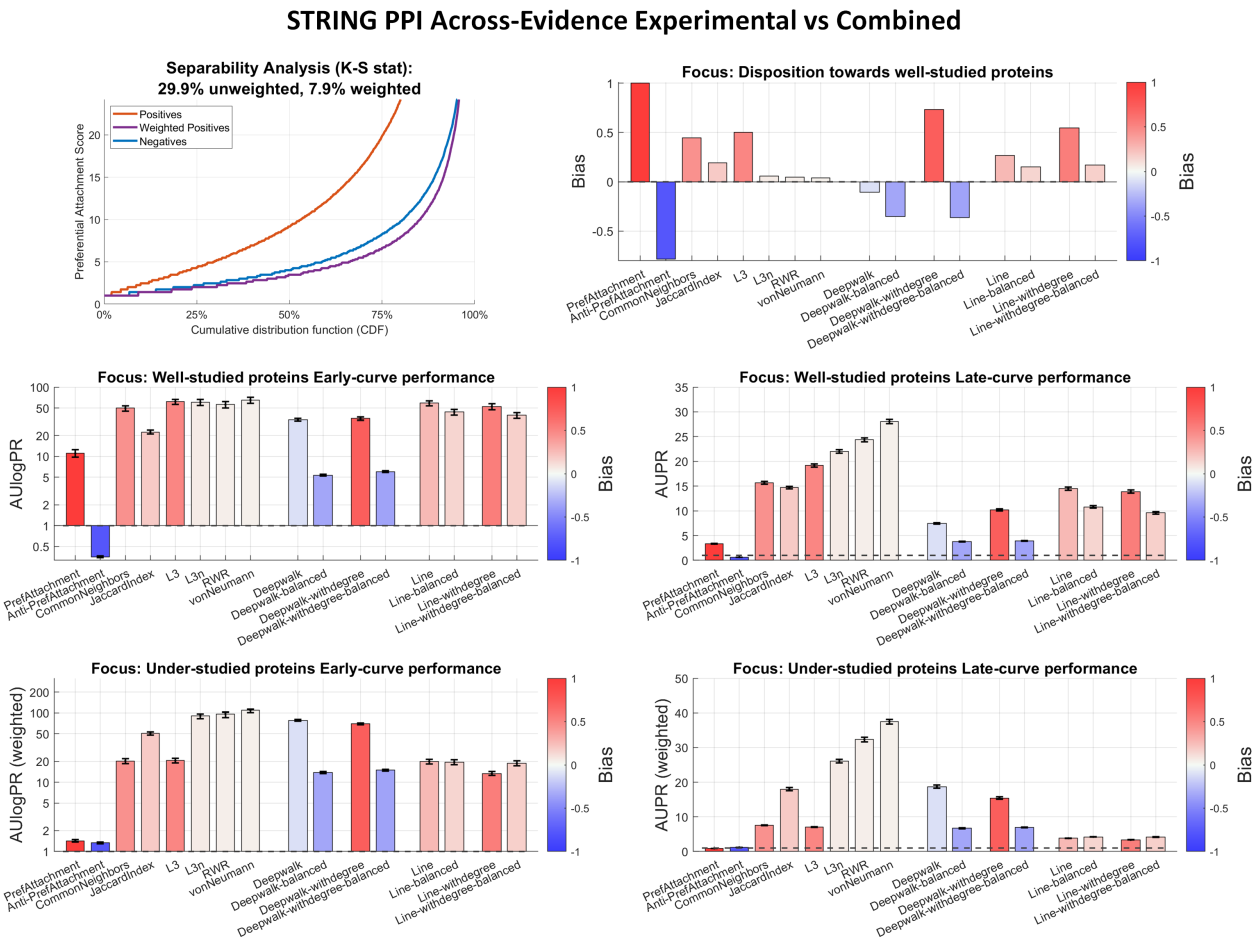

**S. Figure 21: Results of bias-aware evaluation on STRING PPI Across-Evidence (Experimental vs. Combined) setting.** (Top Left) Separability analysis. (Remaining Panels) Evaluation results for each of the five metrics are shown with bar plots: Bias score (based on similarity with preferential attachment), AUPR, AUlogPR, weighted AUPR and weighted AUlogPR to focus on under-studied proteins. In each panel, the coloring is done according to the bias scores.

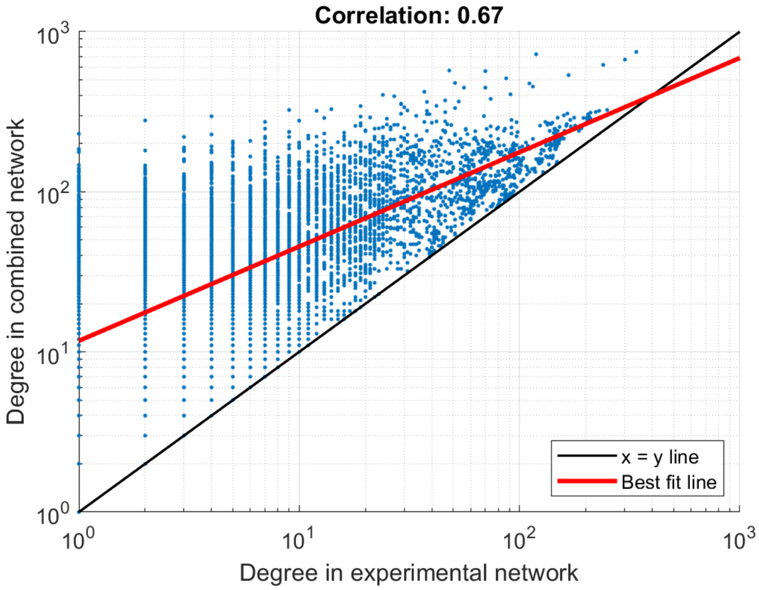

**S. Figure 22: Comparing the node degree distributions in STRING experimental vs. combined networks.**

### A non-exhaustive list of recent publications (2018-2023) regarding link prediction methods developed for biomedical tasks

**[Drug-Disease Associations]**

**[Drug-Drug Interactions]**

**[Transcription Factor Binding - Regulatory Network Inference]**

**[Kinase-Substrate Association]**

**[Protein-Function Prediction]**

**[Protein-Protein Interactions]**

66. Rohit Singh et al. “Topsy-Turvy: integrating a global view into sequence-based PPI prediction”. In: Bioinformatics 38.Supplement 1 (2022), pp. i264–i272.
67. Samuel Sledzieski et al. “D-SCRIPT translates genome to phenome with sequence-based, structure aware, genome-scale predictions of protein-protein interactions”. In: Cell Systems 12.10 (2021), pp. 969–982
68. Paola Paci et al. “Gene co-expression in the interactome: moving from correlation toward causation via an integrated approach to disease module discovery”. In: NPJ systems biology and applications 7.1 (2021), pp. 1–11.
69. Kapil Devkota, James M Murphy, and Lenore J Cowen. “GLIDE: combining local methods and diffusion state embeddings to predict missing interactions in biological networks”. In: Bioinformatics 36.Supplement 1 (2020), pp. I464–i473
70. Kovács, István A., et al. "Network-based prediction of protein interactions." Nature communications 10.1 (2019): 1240.
71. Hu, Lun, et al. "A survey on computational models for predicting protein–protein interactions." Briefings in bioinformatics 22.5 (2021): bbab036.
72. Yuen, Ho Yin, and Jesper Jansson. "Better link prediction for protein-protein interaction networks." 2020 IEEE 20th International Conference on Bioinformatics and Bioengineering (BIBE). IEEE, 2020.
73. Yuen, Ho Yin, and Jesper Jansson. "Normalized L3-based link prediction in protein–protein interaction networks." BMC bioinformatics 24.1 (2023): 59.
74. Chen, Yu, et al. "Protein interface complementarity and gene duplication improve link prediction of protein-protein interaction network." Frontiers in genetics 11 (2020): 291.
75. Wang, Xiaojuan, et al. "Ppisb: A novel network-based algorithm of predicting protein-protein interactions with mixed membership stochastic blockmodel." IEEE/ACM Transactions on Computational Biology and Bioinformatics 20.2 (2022): 1606-1612.
76. Hu, Lun, et al. "A novel network-based algorithm for predicting protein-protein interactions using gene ontology." Frontiers in Microbiology 12 (2021): 735329.
77. Hu, Lun, et al. "A distributed framework for large-scale protein-protein interaction data analysis and prediction using mapreduce." IEEE/CAA Journal of Automatica Sinica 9.1 (2021): 160-172.
78. Yu, Bin, et al. "Prediction of protein–protein interactions based on elastic net and deep forest." Expert Systems with Applications 176 (2021): 114876.
79. Casadio, Rita, Pier Luigi Martelli, and Castrense Savojardo. "Machine learning solutions for predicting protein–protein interactions." Wiley Interdisciplinary Reviews: Computational Molecular Science 12.6 (2022): e1618.
80. Savojardo C, Martelli PL, Casadio R. Protein–protein interaction methods and protein phase separation. Annu Rev Biomed Data Sci. 2020; 3(1): 89–112.
81. Wang L, Wang H-F, Liu S-R, Yan X, Song K-J. Predicting protein-protein interactions from matrix-based protein sequence using convolution neural network and feature-selective rotation Forest. Sci Rep. 2019; 9(1): 9848.
82. Yang F, Fan K, Song D, Lin H. Graph-based prediction of protein-protein interactions with attributed signed graph embedding. BMC Bioinformatics. 2020; 21(1): 323.

**[Protein Interaction Site Prediction]**

**[Miscellaneous, Link Prediction Tasks in Biomedical Context]**
